# Tissue-Specific Transcriptome Responses to Fusarium Head Blight and Fusarium Root Rot

**DOI:** 10.1101/2022.04.07.487462

**Authors:** J F. Haidoulis, P. Nicholson

## Abstract

Fusarium head blight (FHB) and Fusarium root rot (FRR) are important diseases of small-grained cereals caused by *Fusarium* species. While host response to FHB has been subject to extensive study, very little is known about response to FRR and the transcriptome responses of FHB and FRR have not been thoroughly compared. *Brachypodium distachyon* (Bd) is an effective model for investigating host responses to both FHB and FRR. In this study the transcriptome response of Bd to *F. graminearum* (Fg) infection of heads and roots was investigated. An RNA-seq analysis was performed on both Bd FHB and FRR during the early infection. Additionally, an RNA-seq analysis was performed on *in vitro* samples of Fg for comparison with Fg gene expression *in planta*. Differential gene expression and gene-list enrichment analyses were used to compare FHB and FRR transcriptome responses in both Bd and Fg. Differential expression of selected genes was confirmed using RT-qPCR. Most genes associated with receptor signalling, cell-wall modification, oxidative stress metabolism, and cytokinin and auxin biosynthesis and signalling genes were generally upregulated in FHB or were downregulated in FRR. In contrast, Bd genes involved in jasmonic acid and ethylene biosynthesis and signalling, and antimicrobial production were similarly differentially expressed in both tissues in response to infection. A transcriptome analysis of predicted Fg effectors with the same infected material revealed elevated expression of both core tissue-independent genes including cell-wall degradation enzymes and the gene cluster for DON production but also several tissue-dependent genes including those for aurofusarin production and cutin degradation. This evidence suggests that Fg modulates its transcriptome to different tissues of the same host.

## Introduction

*Fusarium graminearum* is a destructive plant pathogen that can cause the economically important disease Fusarium Head blight (FHB) in suitable Poaceae hosts like bread wheat (*Triticum aestivum*), barley (*Hordeum vulgare*), and rye (*Secale cereale*) (Parry et al., 1995). FHB disease is characterised by lesions in the floral bract and caryopsis caused by extensive cell death (Boddu et al., 2006, Lewandowski et al., 2006). Over time discoloration of the peduncle and bleaching of the inflorescence occur and infected florets will produce small and shrivelled kernels due to the destruction of starch and protein (Goswami and Kistler, 2004, Guenther and Trail, 2005). During FHB, *F. graminearum* behaves as a facultative hemibiotroph with a necrotrophic phase preceded by a biotrophic phase (Jansen et al., 2005, Brown et al., 2010). Most wheat plant tissues are susceptible to infection (Miedaner, 1997), and other *Fusarium* diseases include Fusarium Crown Rot (FCR), Fusarium Root Rot (FRR), and seedling blight. FRR causes necrosis of root tissue leading to reduced root, shoot length, biomass, and yield loss (Mergoum et al., 1998, Beccari et al., 2011, Wang et al., 2015). During FRR penetration, sporulation, and necrosis occur rapidly (Wang et al., 2015), and systemic migration of the pathogen via the vascular system can result in FCR and, in extreme cases, to FHB (Beccari et al., 2011, Wang et al., 2015). Due to the inherent difficulty in studying root diseases, much less is known about FRR than FHB.

Apart from affecting grain yield and quality of the cereal crop, *F. graminearum* can synthesise deoxynivalenol (DON), and nivalenol (NIV) mycotoxins that contaminate grain and pose a risk to human and animal consumers (Antonissen et al., 2014, Payros et al., 2016).

*Fusarium graminearum* is known to utilise a combination of cell wall-degrading enzymes (CWDEs) and DON to overcome host defences in small-grain cereals. Infection is also accompanied by an increase in secreted effectors, changes to pathogen molecular transport and signalling, and changes to secondary metabolite and nutrient metabolism (Kikot et al., 2009, Brown et al., 2017, Lysøe et al., 2011). Cereal hosts have been shown to respond to FHB through the deployment of numerous defences including deoxynivalenol (DON) detoxication, host metabolism changes, cell wall development changes, and synthesis of antimicrobial compounds and *PATHOGENESIS-RELATED* (*PR*) proteins (Boddu et al., 2006, Jia et al., 2009, Pasquet et al., 2014).

Transcriptomics is an effective tool to investigate molecular responses during pathogenesis. Several studies have investigated *F. graminearum*-induced changes in gene transcription during infection of wheat and barley (*Hordeum vulgare*) using methods such as microarray analysis and real-time quantitative PCR (RT-qPCR) (Boddu et al., 2006, Jia et al., 2009, Li and Yen, 2008). More recently, RNA-seq has been utilised to study wheat responses to FHB and FCR (Powell et al., 2017a, Pan et al., 2018, Wang et al., 2018a). Likewise, several studies have investigated the transcriptome of *F. graminearum* during Fusarium head blight (FHB) disease of small grain cereals, in most cases through microarray analysis (Lysøe et al., 2011, Harris et al., 2016, Brown et al., 2017). RNA-seq technology has been used to-date, in terms of *F. graminearum* disease transcriptomics, for FRR (Ding et al., 2022) and FHB (Pan et al., 2018).

Of the canonical defence phytohormones, the consensus is that salicylic acid (SA) - regulated responses are associated with resistance to biotrophic pathogens whereas jasmonic acid (JA) and ethylene associated responses are linked to resistance to necrotrophic pathogens (Pieterse et al., 2012, Glazebrook, 2005, Bari and Jones, 2009). As a result of the hemibiotrophic lifestyle of *F. graminearum*, SA and JA/ethylene pathways have both been shown to be important for defence (Makandar et al., 2011, Ding et al., 2011, Wang et al., 2018a). Many other phytohormones including abscisic acid (ABA), gibberellic acid (GA), auxin, and cytokinin have also been implicated in response to *F. graminearum* infection (Pan et al., 2018, Wang et al., 2018a, Powell et al., 2017a). In a previous study we showed that exogenous application of many phytohormones induced significant effects on FHB and FRR resistance (Haidoulis and Nicholson, 2020). However the phytohormones SA, JA, and ethylene induced opposing effects on resistance to FHB and FRR whereas cytokinin and auxin induced similar effects in the two tissues.

*Brachypodium distachyon* (purple false brome) is a temperate monocotyledonous plant in the Pooideae sub-family. Similar to wheat, both *B. distachyon* roots and florets can be infected by *F. graminearum* infection which permits the investigation of both FHB and FRR in this host (Peraldi et al., 2011, Pasquet et al., 2014).

It is unclear whether *F. graminearum* differentially expresses genes and deploys tissue- specific effectors within the same host. The aim of the present study is to compare the transcriptomes of both host (*B. distachyon*) and pathogen (*F. graminearum*) during early FHB and FRR infection to identify similarities and differences related to the host tissue. We believe this to be the first study to examine both host and pathogen gene expression within different tissues in the same host. The data reveals that gene expression of *B. distachyon* defence and *F. graminearum* virulence occur in both a tissue-specific manner and non-specific manner suggesting that there are core processes alongside bespoke tissue-related responses in both host and pathogen.

## Materials and Methods

### Plant and Fungal Material and Growth Conditions

The *B. distachyon* accession Bd3-1 was obtained from the John Innes Centre, Norwich, UK. *B. distachyon* seed preparation and growth conditions were the same as in (Haidoulis and Nicholson, 2020) except for a three day seed stratification for root assay.

The *F. graminearum* isolate PH1 was obtained from the John Innes Centre, Norwich, UK and used for all experiments. For the FRR assay mycelium inoculum and FHB conidial suspension, *F. graminearum* was prepared as in (Haidoulis and Nicholson, 2020). The *F. graminearum in vitro* control samples (1 x 10^4^ conidia/ml) were grown in sterile Czapek-Dox liquid medium with 1 unit/ml penicillin-streptomycin for four days and incubated in a shaker at 25°C and 200 rpm.

### Sample Inoculation and Preparation

For FRR assays, ten stratified seeds were placed on 9 cm^2^ filter paper square on a 50 ml 0.8% agar (Fischer Science) in square Petri-dishes. All plates were angled at 70° from the horizontal in a plant propagator tray and incubated for three days at 22°C (16h/8h - light/dark photoperiod) before inoculation. Mycelial slurry was prepared from blended seven-day old mycelium from a PDA petri-dish with 1 ml sterile deionised water. After three days, Bd3-1 roots were inoculated at three positions per root with a mycelial PDA slurry or PDA slurry using a 10 ml syringe. After one day, the inoculum slurry was removed, and roots were rinsed with water. For each biological replicate, ten roots were cut and frozen in liquid nitrogen.

For FHB assays, Bd3-1 spikelets were sprayed at mid-anthesis with conidial inoculum (1 x 10^6^ conidia/ml) in sterile distilled water before the dark photoperiod. Both inoculum suspension and water control were amended with 0.05% Tween 20. Plants were incubated at 22°C for 3 d at high humidity. After three days, for each biological replicate, three infected and mock inoculated spikelets from randomly selected plants were cut and immediately frozen in liquid nitrogen.

RNA was extracted from both spike and root tissues using QIAGEN RNAeasy kit as per the manufacturers protocol. RNA was then cleaned using Turbo DNA-free kits as per standard protocol with two rounds of Turbo DNAse treatment. RNA samples were quantified and quality checked using a Qubit and TapeStation (performed by Genewiz).

### Library Preparation RNA-seq and Analysis

Library preparation was performed at Genewiz and sequenced using Illumina HiSeq, PE 2x150bp sequencing configuration with a single index per lane. RNA-seq illumina reads FASTA data, obtained from Genewiz, were analysed on the Galaxy platform (Afgan et al., 2016). FastQC (G.V.0.72) was used for sample FASTA reads as a quality check. Using Trimmomatic (G.V.0.36.5), paired-end FASTA reads were trimmed with the settings: ‘Sliding window’ (4 bases), ‘leading’ and ‘trailing’ ends (3 bases each), and TrueSeq3 illumina clip was used to remove illumina adaptor sequences. Trimmed FASTA reads were quality checked again with FASTQC (G.V.0.72). Trimmed FASTA reads were aligned to the most recent version of the Bd assembly (Bd21 JGI v3.0)(Phytozome JGI V12.1.5, (Initiative, 2010, Goodstein et al., 2012)) using HISAT (G.V.2.10). Gene annotations were assigned using Stringtie v3.1 (G.V.1.3.4) with annotations (Phytozome JGI, *B. distachyon* v3.1 (Initiative, 2010, Goodstein et al., 2012)). Stringtie gene counts for FHB and FRR were differentially compared to respective control samples with DEseq2 (G.V.2.11.40.2).

The predicted functions for *B. distachyon* genes were obtained from Ensembl Genomes (Howe et al., 2020), UniProt (Consortium, 2018), BrachyPan (Goodstein et al., 2012), (Gordon et al., 2017), *B. distachyon* v3.1 from Phytozome JGI (V12.1.5) (Initiative, 2010, Goodstein et al., 2012), or (Kakei et al., 2015, Kouzai et al., 2016). The predicted gene functions within the heatmaps with a prefix and percentage homology denotes the percentage of *B. distachyon* sequence that matches the orthologous sequence obtained from Ensembl Genomes (Howe et al., 2020)), were derived from the Arabidopsis Information Resource (TAIR10) database (Berardini et al., 2015), or from previous studies (Yazaki et al., 2004, Jain et al., 2006b, Jain et al., 2006a, Tsai et al., 2012, Jain et al., 2005). Then, if necessary, the *B. distachyon* homologue/s were identified within the Ensembl Genomes database (Howe et al., 2020) (At: *thaliana* TAIR10, Os: *O. sativa* RGSP-1.0, Hv: *H. vulgare* IBSC_v2, or Zm: *Z. mays* B73_RefGen_v4)) and were then searched for within the RNA-seq dataset.

The same pipeline described above was used for *F. graminearum* reads with the same samples but aligned to the *F. graminearum* PH1 genome assembly and gene annotation (European Nucleotide Archive; GCA_900044135.1, study PRJEB5475 (King et al., 2015)). FHB and FRR sample gene counts were separately compared against the same *F. graminearum in vitro* control samples using DEseq2. The average of normalised gene counts per biological replicate from FHB and FRR DEseq2 is presented. Differentially expressed *F. graminearum* genes in FHB and FRR were filtered for potential effectors using the PH1 v5.0 secretome prediction script (Brown et al., 2012). The online databases Ensembl Genomes (Howe et al., 2020), UniProt (Consortium, 2018), and protein sequence BLAST (Sayers et al., 2020) were used to predict *F. graminearum* gene functions.

### Time-course RT-qPCR

FHB and FRR samples were prepared as described before. Samples from different plants were harvested at 3 dpa, 5 dpa, and 7 dpa for FHB and 1 dpa, 3 dpa, and 5 dpa for FRR, with at least three biological replicates per treatment and time-point. For *F. graminearum* genes, only one time point was used for the *in vitro* control which was subsequently compared to all three infected FHB and FRR time-point samples. After sample RNA extraction using QIAGEN RNAeasy kit as per the manufacturers protocol and DNase treatment as described before, first strand synthesis of RNA was performed with Invitrogen SuperScript III Reverse Transcriptase (Invitrogen) as per the manufacturers protocol. Reverse transcriptase qPCR was performed with 2 µl cDNA, 5 µl of 2x SYBR Green JumpStart Taq ReadyMix (Sigma-Aldrich), 0.6 µl of 10 µM for each primer (Supplementary Table S1), in a final volume of 10 µl. RT-qPCR reactions were prepared in a Framestar-480/384 well plate with BioRAD microseal B adheasive film. Thermocycling was carried out on a Roche LightCycler LC480 on SYBR green 1 scan mode with the following parameters: 300 s 95°C, 45 x (94°C 10 s, 58°C (or 60°C for Bradi1g57590) 10 s, 72°C 10 s, 75°C 2 s (single acquisition)) and subsequently analysis of dissociation by ramping 95°C 5s, 60°C 60 s, 97°C (continuous) followed by 40°C for 30 s. LC480 raw data was converted with LC480 conversion software and analysed for primer efficiency and Cq values with the LinRegPCR tool (Ruijter et al., 2009). Log fold changes were calculated from Cq values using the following equations: Δ Cq = (Gene of interest Cq – housekeeping gene Cq), ΔΔ Cq = (Infected treatment Δ Cq – mock treatment Δ Cq), Log2 fold change = Log2 (Primer efficiency^ΔΔ Cq^) (Pfaffl, 2001).

All primers (Sigma Aldrich UK. Primers) for gene targets (Supplementary Table S1), unless otherwise stated, were designed using Primer 3 (Kõressaar et al., 2018, Untergasser et al., 2012, Koressaar and Remm, 2007) on a single CDS exonic region and avoiding untranslated (UTR) regions. The best housekeeping gene *GAPDH* (Supplementary Table S1) for these samples was experimentally determined and analysed on NormFinder in GenEx V6 using cDNA obtained from both control and infected root and spike material.

### Statistics and Software

The RNA-seq p-value and p-adj values were outputs from Galaxy DEseq2. A standard Student’s t-test on Microsoft Excel was used for time-course RT-qPCR Cq data. Heatmaps were prepared on R studio using ‘pheatmap’ and ‘rcolorbrewer’ package. The normalised transcript counts (reads) were transformed (Log2 (x + 1)) and then scaled per gene (row). Hierarchical clustering of genes (rows) used Euclidean distance metric with complete-linkage clustering. Volcano plots were prepared on R studio using the packages ‘ggplot2’, ‘ggrepel1’, and ‘EnhancedVolcano’ (Blighe et al., 2019). Venny V2.1 was used for Venn Diagrams and sorting treatment groups (Oliveros, 2018). Graphs for time-course experiment were prepared on GraphPad Prism V5. Gene ontology enrichment for *B. distachyon* was achieved using The Gene Ontology Resource (Database released 2021-08-18) and PANTHER Overrepresentation Test (Released 20210224) (Ashburner et al., 2000, 2021, Mi et al., 2019, Mi et al., 2021). Gene Ontology enrichment plots were produced as described in (Bonnot et al., 2019, Supek et al., 2011). Fisher’s exact test type with FDR correction was used for GO enrichment and default REVIGO settings were used. Gene-list enrichment for *Fusarium graminearum* was performed using KOBAS 3.0 (Wu et al., 2006, Xie et al., 2011, Ai and Kong, 2018).

## Results

### Fusarium Head Blight and Fusarium Root Rot display distinct global transcriptome responses to infection

The time points 3 days-post inoculation (dpi) for FHB and 1 dpi for FRR represent the earliest stage at which symptoms were visible for the two diseases. Differential gene expression analysis was performed on total gene counts of diseased *B. distachyon* floral and root tissues in comparison to respective mock-inoculated treatments. Coverage to the *B. distachyon* (accession Bd21) assembly was between 80-95% coverage. With no log-fold change threshold, there were 6,158 genes significantly differentially expressed in response to FHB (p-adj < 0.05) (Figure 1A), whereas 8,568 genes were significantly differentially expressed in response to FRR (p-adj < 0.05) (Figure 1B). Approximately 17% of the genes significantly differentially expressed in response to FRR exceeded the 2 Log-fold change threshold, whereas 29% of the genes significantly differentially expressed in response to FHB exceeded the 2 Log-fold change threshold.

**Figure 1:**
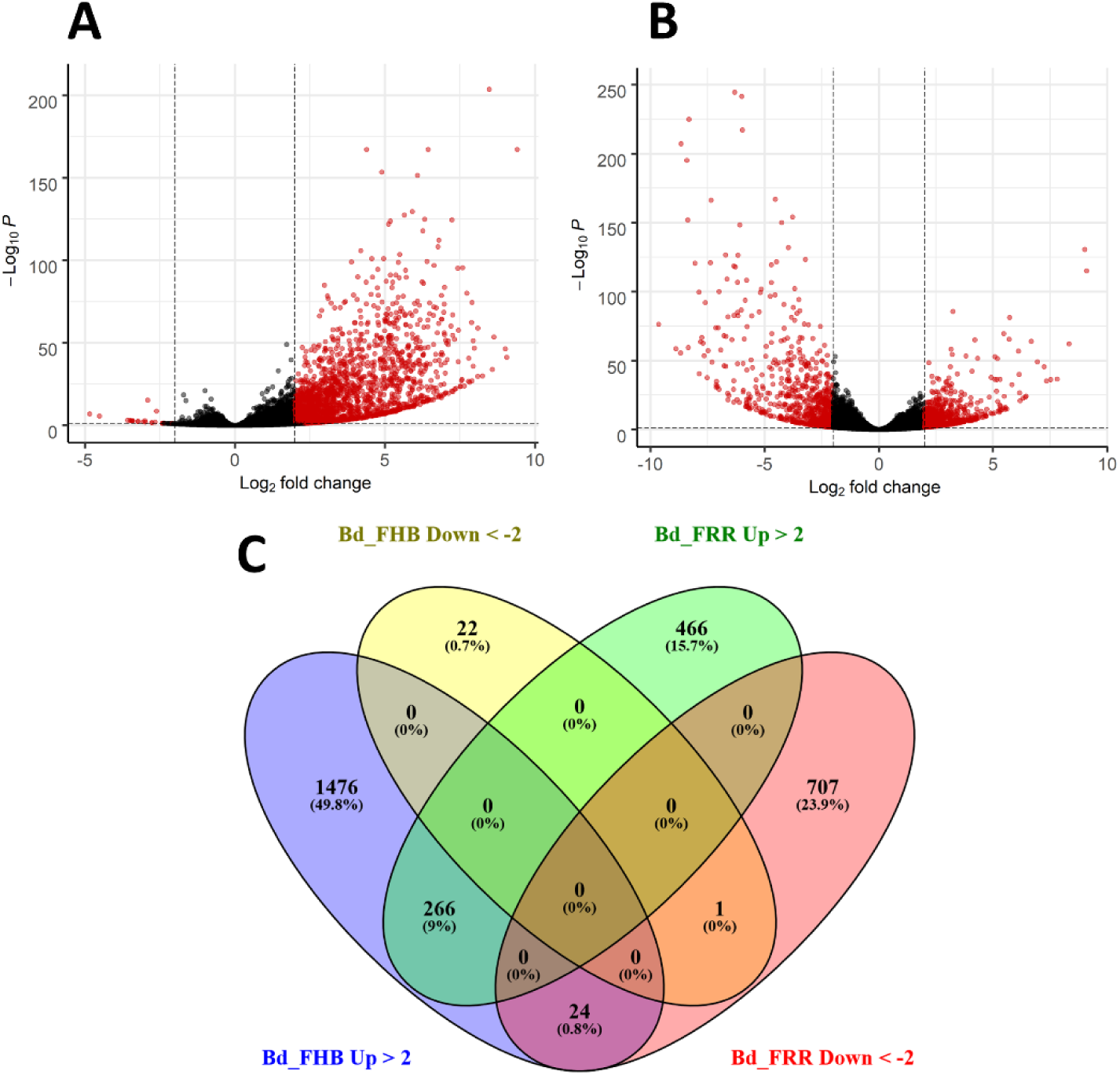
Summary of expression of *B. distachyon* genes in response to FHB and FRR. (**A**) Differential gene expression in response to FHB. (**B**) Differential gene expression in response to FRR. (**A** and **B**) Each dot represents a gene (Supplementary Table S3), excluding those without a p-adj value. The y- axis is the -Log10 of the p-adj value (p-adj < 0.05) denoted by a dotted line. The x-axis is the Log2 fold change with a cut-off of 2 denoted by two dotted lines. Thus a red dot denotes that a gene was statistically significantly differentially expressed. (**C**) A comparison of genes between FHB and FRR datasets. The threshold of -2 ≤ x ≥ 2 Log-fold change and p-adj < 0.05 was applied to all genes. Abbreviations: Bd (*B. distachyon*), Up (Upregulated), Down (Downregulated), FRR (Fusarium Root Rot), FHB (Fusarium Head Blight).

The transcriptome response between FHB and FRR was compared (Figure 1C). Following FHB infection the number of upregulated genes (1766) was much greater than the number of downregulated genes (23) (Figure 1A). In contrast, similar numbers of genes were upregulated and downregulated in response to FRR (Figure 1B). There were relatively few genes that were upregulated (226) or downregulated (1) in response to both FHB and FRR. There were more genes exclusively upregulated (466 genes) and downregulated (707 genes) in response to FRR but the most pronounced difference was observed for FHB where 1,476 genes were exclusively upregulated. In contrast, only 22 genes were exclusively and significantly downregulated in response to FHB. A small group of 24 genes were upregulated in response to FHB and downregulated in response to FRR (Figure 1C). Of these 24 (Supplementary Table S4), notable genes include a xyloglucan endotransglucosylase (Bradi3g31767), a pathogenesis-related protein 1 (Bradi3g53681), a disease resistance protein RPP13-related (Bradi1g29381), a peroxidase (Bradi5g27150), an endoglucanase (Bradi3g36210), a RING-type E3 ubiquitin transferase (Bradi3g52120), and an expansin (Bradi3g09960).

### Gene Ontology Enrichment Analysis Reveals Differentially Expressed Pathways in *B. distachyon*

Gene Ontology (GO) enrichment analysis was performed on the different treatment groups: FHB upregulated, FRR upregulated, and FRR downregulated (Figure 2). A total of 67 GO biological processes were significantly overrepresented in one or more of the treatment groups (Figure 2). Most GO terms were overrepresented in a tissue-specific manner. For overrepresented FHB-specific processes, notable GO terms were receptor signalling (GO:0007178, GO:0007167, GO:0007166), defence/immune response and response to microorganisms (GO:0009611, GO:0031347, GO:0002376, GO:0042742, GO:0002239, GO:0002229, GO:0009607, GO:0044419, GO:0006950), Glutathione metabolic process (GO:000674), indole compound biosynthesis and metabolism (GO:0042435, GO:0042430), jasmonic acid signalling (GO:2000022), oxylipin biosynthesis and metabolism (GO:0031408, GO:0031407), phenylpropanoid metabolism (GO:0009698), and phosphorylation-associated processes (GO:0000160, GO:0006468, GO:0016310, GO:0006793) (Figure 2). The pathways for response to stimulus and stress (GO:0050896, GO:0006950), and phosphorylation-associated processes (GO:0006468, GO:0016310, GO:0006793) showed the greatest number of significantly expressed genes expected in the pathway for the GO analysis (Figure 2). There were insufficient genes and no significant biological processes for the FHB-downregulated gene list. The GO pathways: cellular response to chemical stimulus (GO:0070887) and tryptophan biosynthesis (GO:0000162), were enriched exclusively in FRR upregulated genes (Figure 2). Only three GO processes were similarly overrepresented within the two tissues from upregulated gene: response to ethylene (GO:0009723), the ethylene-activated signalling pathway (GO:0009873) and aromatic acid family metabolic process (GO:0009073) (Figure 2). There were far more GO processes for downregulated FRR differentially expressed genes than upregulated genes, notably an overrepresentation in cellular oxidant detoxification (GO:0098869), cell wall modification (GO:0071669, GO:0046274, GO:0071554, GO:0006073, GO:0045229), fluid and water transport (GO:0042044, GO:0006833), response to toxic substance (GO:0009636), and carbohydrate metabolism (GO:0006073, GO:0044262, GO:0005976, GO:0005975) (Figure 2). The pathway biological process (GO:0008150) showed the greatest number of significantly expressed genes expected in the pathway for the GO analysis (Figure 2). Three GO processes for response to oxidative stress (GO:0006979), ROS metabolism (GO:0072593), and hydrogen peroxide detoxification (GO:0042744) were overrepresented in both upregulated and downregulated FRR-differentially expressed genes. These were among the most significant and greatest fold enrichment for downregulated processes (Figure 2).

**Figure 2:**
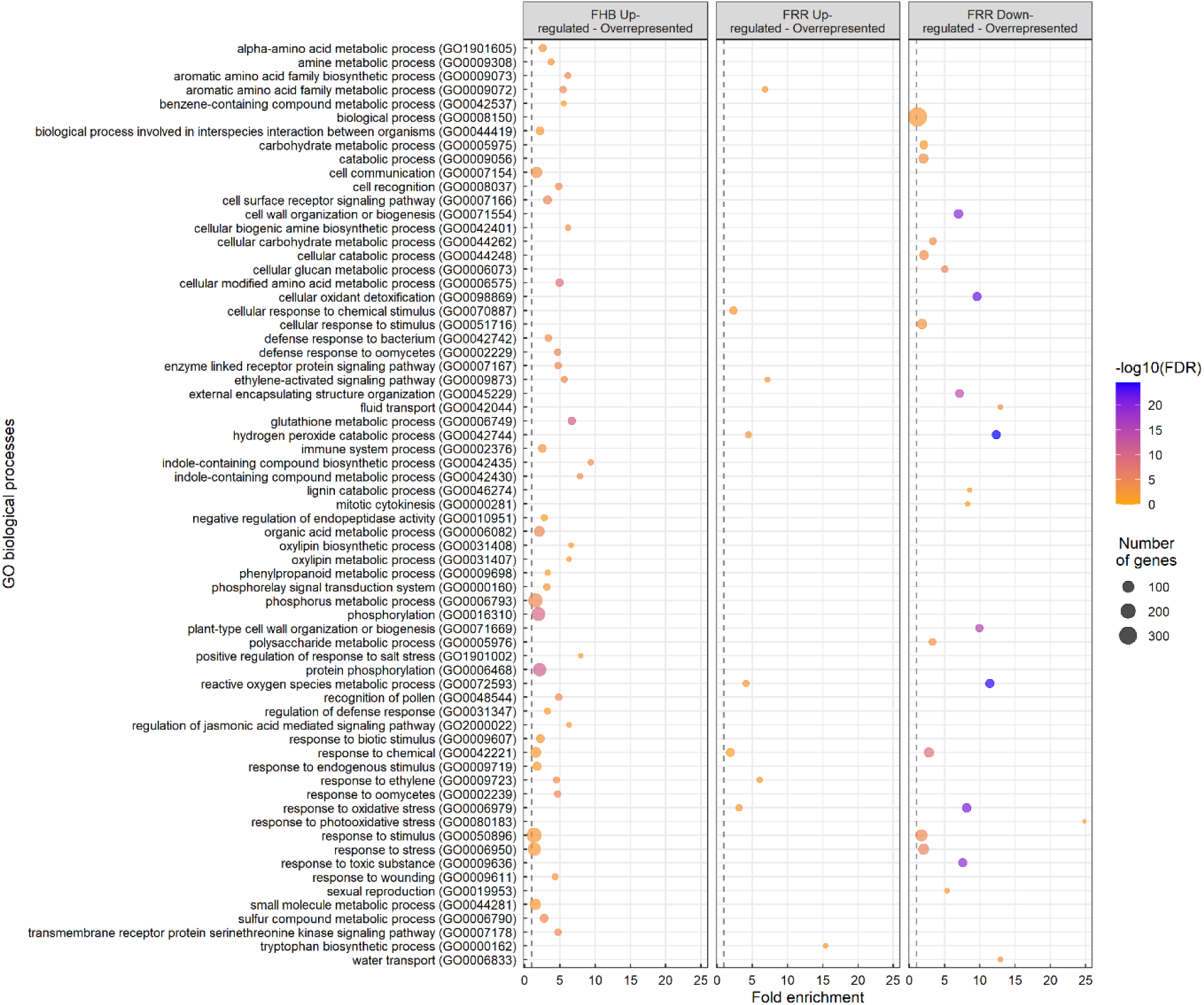
Overrepresented biological processes from significantly upregulated and downregulated genes in FHB and FRR. The dot size represents the number of genes within the process and the dot colour represents the enrichment significance (-log10 (False Discovery Rate (FDR) - corrected p-value)). The vertical dotted lines denote a fold enrichment threshold of 1 signifying that the biological processes are overrepresented. Data in Supplementary Table S5.

### Several gene groups were differentially expressed in response to FHB and FRR

Significantly upregulated or downregulated *B. distachyon* genes (p-adj < 0.05) were grouped based on predicted function and compared between FHB and FRR. Differential expression was defined as having a log-fold threshold more than 2-fold or less than -2-fold change, and a p-adjusted value of less than 5% significance level (p < 0.05). Several genes were found to encode similar products and function within the same role. These were grouped and described in the following sub-sections, focusing on pathways that have roles in plant-pathogen interactions and defence.

### Phytohormone-related genes

Phytohormones play important roles in Fusarium resistance. Exogenous application of JA, ethylene, auxin, and cytokinin was shown to significantly affect resistance to both FHB and FRR in *B. distachyon* (Haidoulis and Nicholson, 2020). Here we add to these results and show that phytohormone-related changes in response to infection also occur at the transcriptional level. In a similar fashion, based on predicted-function of genes, the phytohormones JA, ethylene, auxin, cytokinins, and to a lesser extent (ABA and SA) were the main phytohormones significantly altered in transcription to FHB and FRR in the RNA-seq data (Supplementary Table S6). The transcription of genes related to phytohormones SA, auxin, cytokinin, and ABA were mostly differentially expressed in FHB and FRR. Most SA and ABA – related genes were upregulated in FHB. Most auxin and cytokinin – related genes were downregulated in FRR and were either upregulated or not expressed in FHB. In contrast, ethylene, and to a lesser extent JA-related, transcription tended to be similar in FHB and FRR.

There were 20 jasmonic acid-related genes differentially expressed between FHB and FRR (Figure 3A). Only the JA-repressor *JAZ* genes (Bradi3g23190, Bradi4g31240), and the JA biosynthesis lipoxygenase *LOX2* gene Bradi3g39980 were upregulated in both FHB and FRR (Figure 3A) while the remaining five *JAZ* genes increased in expression only in FHB (Figure 3A). Furthermore, all the biosynthetic *OPR* genes (Bradi1g05870, and Bradi1g05860), lipoxygenase genes, and *4CL* were exclusively upregulated in FHB (Figure 3A). In contrast, the lipoxygenase genes Bradi1g09260 and Bradi1g11680, and Jasmonate O-methyltransferase (Bradi1g43080), were downregulated in response to FRR (Figure 3A). Overall, JA-biosynthetic and signalling genes appeared primarily FHB-responsive. Interestingly there were differences in basal expression of several JA-related genes in the two tissues. The genes Bradi3g43920, Brad1g72610, and Bradi4g20220 had much higher transcript counts in non-inoculated spikes as opposed to roots while all *OPR* genes, including Bradi1g09260, Bradi5g08650, Bradi5g11590, and Bradi3g37300 had much higher transcript counts in non-inoculated roots as opposed to spikes (Figure 3A).

**Figure 3:**
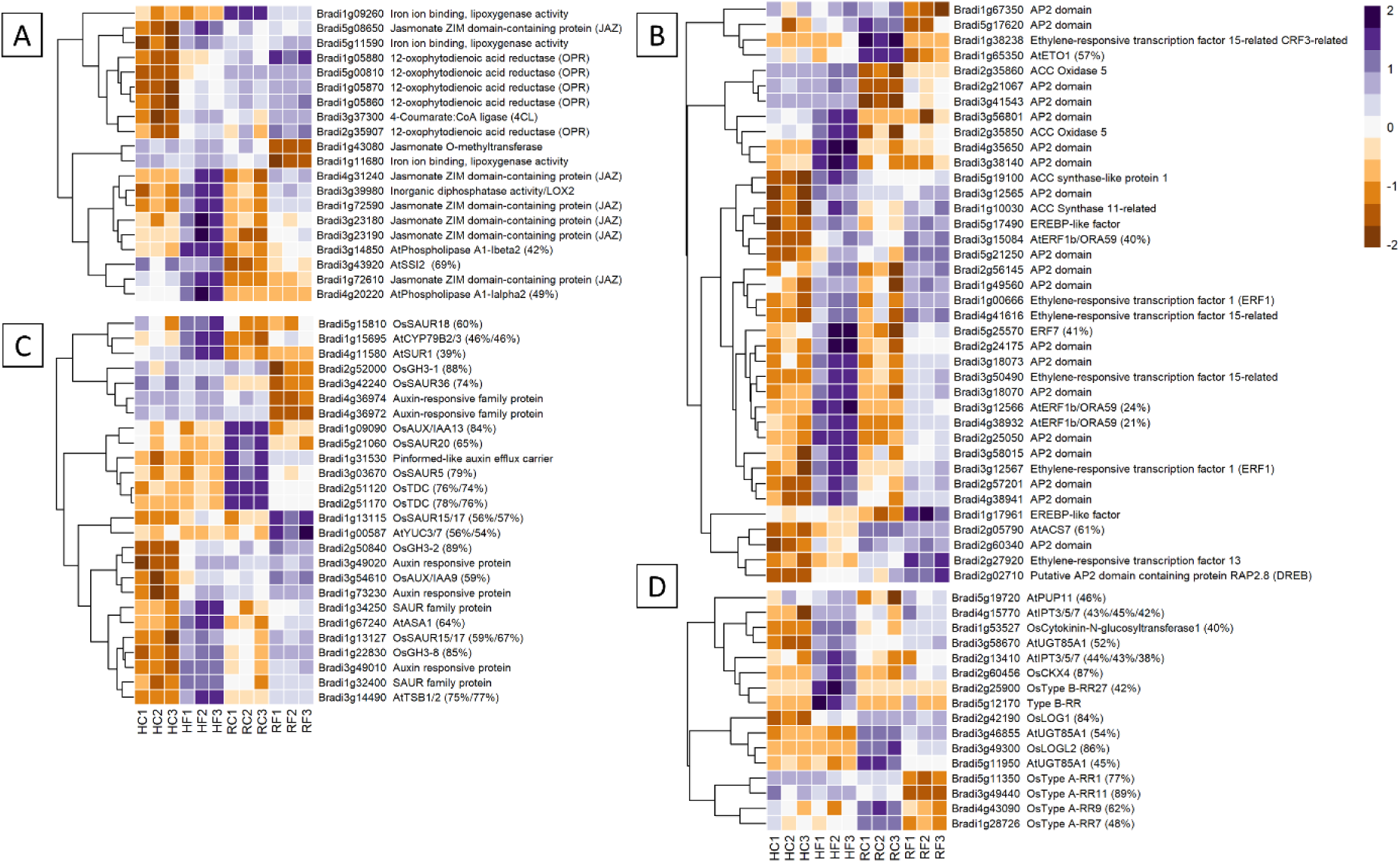
The most expressed or repressed phytohormone-associated *B. distachyon* genes. JA (**A**), ethylene (**B**), auxin (**C**), and cytokinin (**D**) related genes are displayed. These genes showed a log-fold change of -2 ≤ Log2 ≥ 2 with a p-adj < 0.05 in response to either FHB, FRR, or both. The scale bar on the right is the Z-score for all the heatmaps. Some gene functions have a prefix denoting that they were derived from *A. thaliana* (At) or *O. sativa* (Os) and they include the percentage of *B. distachyon* sequence that matches the homologous sequence. Three biological replicates for each of the four treatments are displayed as columns abbreviated as HC (Head-FHB control), HF (Head-FHB fungus), RC (Root-FRR Control), RF (Root-FRR fungus). The control samples (HC1-HC3, RC1-RC3) are normalised transcript counts from mock inoculated head (water with Tween 20) and root (PDA slurry) *B. distachyon* tissues and were separately analysed through the RNA-seq pipeline with the respective inoculated sample tissues.

Ethylene-associated genes were the largest phytohormone-related group expressed for FHB and FRR (38 genes) (Figure 3B). As for JA, most of the ethylene biosynthesis genes encoding *ACS* and *ACO* genes were upregulated in response to FHB (Bradi5g19100, Bradi2g35850, Bradi1g10030, Bradi2g05790). The biosynthesis regulator Bradi1g65350 (orthologues to *AtETO1*) was exclusively downregulated in response to FRR. Conversely, *ACO5* (Bradi2g35860) was upregulated in response to FRR. Differential expression of many ethylene-associated genes was just below the threshold (Log-fold change > 2) for FRR (Supplementary Table S6). Most ethylene-associated genes encoded well known downstream transcription factors such as *ERF1/ORA59* or other less characterised genes encoding proteins with AP2/EREBP domains (Broekaert et al., 2006). The majority showed similar expression patterns in response to FHB and FRR including *ERF1* (Bradi3g12567) and *ERF1b/ORA59* (Bradi4g38932). The genes *ACS7* (Bradi2g05790), *ERF15* (Bradi1g38238), and *ETO1* (Bradi1g65350) had substantially higher basal expression in non-inoculated root tissues compared to non-inoculated spike tissues.

Expression of many of the 26 auxin-associated genes identified, increased in FHB but the response in FRR was mixed (Figure 3C). Auxin biosynthetic orthologues were mainly upregulated in FHB (*AtSUR1* Bradi4g11580, *AtTSB1/2* Bradi3g14490). However, while some were also upregulated (*AtASA1* Bradi1g67240, *AtYUC3/7* Bradi1g00587, *AtCYP79B2/3* Bradi1g15695) others were downregulated in FRR (*OsTDC*, Bradi2g51120 Bradi2g51170). The biggest differences in response between FHB and FRR were for genes downstream of auxin biosynthesis. *GH3* orthologues with known roles in resistance (Ding et al., 2008, Fu et al., 2011, Domingo et al., 2009) were differentially expressed with *OsGH3-8* (Bradi1g22830) upregulated in both FHB and FRR, *OsGH3-2* (Bradi2g50840) expressed in FHB only, and *OsGH3-1* (Bradi2g52000) downregulated only in FRR. The auxin-response factor (*ARF*) genes Bradi3g49010, Bradi3g49020, and Bradi1g73230 were upregulated in FHB whereas Bradi4g36974 and Bradi4g36972 were downregulated in FRR. The response of *SAUR* genes was also highly variable. Bradi1g13127 was upregulated in both FHB and FRR, Bradi5g15810 was upregulated only in FHB, while Bradi5g21060 and Bradi3g42240 were downregulated only in FRR. Lastly, only one auxin transport gene was repressed (Pin-efflux carrier, Bradi1g31530) in FRR.

Rice orthologues were also used to identify cytokinin associated genes in *B. distachyon* (Tsai et al., 2012). From the total of 16 cytokinin-associated genes identified, almost all biosynthetic genes were upregulated in FHB (e.g. *AtUGT85A1* Bradi3g58670, *AtIPT3/5/7* Bradi2g13410, *AtIPT3/5/7* Bradi4g15770, and *OsLOG1* Bradi2g42190). In contrast many biosynthetic genes were downregulated in response to FRR (*OsLOGL2* Bradi3g49300, *AtUGT85A1* Bradi3g46855, *AtUGT85A1* Bradi5g11950). The same was true for downstream signalling Response Regulator (RR) genes. All *OsType B RR* homologues were upregulated in FHB (Bradi2g25900, Bradi5g12170), whereas all *OsType A RR* homologue were significantly downregulated in FRR (Bradi4g43090, Bradi5g11350, Bradi3g49440, Bradi1g28726).

Salicylic acid (SA) is an important phytohormone involved in resistance to biotrophs and generally functions antagonistically to JA (Pieterse et al., 2012, Glazebrook, 2005, Bari and Jones, 2009). The Bd homologues of SA-responsive genes *OsNPR4* (Bradi2g54340), *NPR1 interacting* (Bradi2g27670), *AtCBP60g* (Bradi4g05360), *AtSARD1* (Bradi2g02310), *AtGRX480* genes (Bradi2g08400 and Bradi2g46093) were exclusively upregulated in response to FHB (Supplementary Table S6). Likewise, several systemic acquired resistance (SAR)-associated genes *MES1*-encoding genes were upregulated in FHB (Bradi2g41070, Bradi3g44867, Bradi4g35382) whereas two were downregulated in FRR (Bradi2g52110 and Bradi4g09007). Bradi1g71530, orthologous to *AtALD1*, was also exclusively upregulated in FHB. Similarly, Bradi3g43920 (orthologous to *AtSSI2* involved in SA and JA antagonism (Pieterse et al., 2012)) was upregulated exclusively in response to FRR (Figure 3A). Relatively few SA-related genes were significantly differentially expressed between FHB and FRR compared to the previous four phytohormones, but for the few that were, they were primarily upregulated in FHB as opposed to FRR.

Several ABA-associated genes were also identified as differentially expressed (Supplementary Table S6). The biosynthetic genes homologous to *Arabidopsis* genes *AtNCED2/3/5/9* were generally upregulated in response to FHB (e.g. Bradi1g13760, Bradi1g51850, Bradi1g58580) whereas Bradi1g52740 (orthologous to *AtAAO1/2/3/4*) was downregulated in response to FRR. The homologue of the signalling gene *AtAP2C1* was also upregulated in response to FHB. In contrast, downstream Bradi2g43056, Bradi2g60441, Bradi2g60490 (orthologous to homeostasis *AtBG1* genes) were upregulated in response to both FHB and FRR. Overall, there was a substantial differential in expression of ABA biosynthesis and signalling genes between FHB and FRR. Very few GA and BR-associated genes were identified as being differentially expressed in response to either FHB or FRR (Supplementary Table S6).

### Genes encoding pathogen sensing proteins

Receptor-related genes displayed one of the largest differences in expression between FHB and FRR. The majority were upregulated in FHB but either not expressed or downregulated in FRR. Those that were also upregulated in FRR did so at a much-reduced level relative to FHB (Figure 4A). Putative leucine rich receptor-like kinases (LRR-RLKs) (e.g. Bradi1g22650, Bradi4g11740, and Bradi2g19380), LRR-containing genes (e.g. Bradi3g55006, Bradi4g42828, and Bradi4g42839), and nucleotide binding leucine rich repeat (NBS-LRR) proteins (e.g. Bradi5g15560, Bradi1g56690, Bradi2g21360) were almost all exclusively expressed in FHB (Supplementary Table S7). Only the LRR-RLK Bradi1g32160 was significantly upregulated in FRR (Logfold > 2). The remaining LRR-RLKs were either not expressed or were downregulated in FRR (e.g. Bradi1g75430). LRR-RLKs can be grouped into specific classes (Liu et al., 2017, Rameneni et al., 2015). The LRR-RLK classes 8, 10b, 12, 14, and 15 were only expressed in FHB (except Bradi1g32160) (Figure 4A and Table S7). On the other hand, LRR-RLK classes 7a, and 11 were exclusively downregulated in FRR. NBS-LRR-encoding genes were primarily upregulated in FHB and downregulated in response to FRR except for Bradi2g18840 which was highly expressed in both tissues (Figure 4A and Table S5). Disease resistance rpp13- like proteins (Bradi1g29381, Bradi5g01167, Bradi4g21942 Bradi4g09247) showed different differential expression between FHB and FRR (Figure 4A and Supplementary Table S7). Many genes encoding wall-associated receptor kinase (WAK) galacturonan-binding genes (e.g. Bradi2g02440, Bradi2g02450, Bradi2g02537), WAK receptor-like proteins (e.g. Bradi2g17520, Bradi3g01170, Bradi3g39670), or WAK receptor-like protein kinases (e.g. Bradi3g39670, Bradi2g02470, Bradi2g17520) were exclusively upregulated in FHB (Supplementary Table S7).

**Figure 4:**
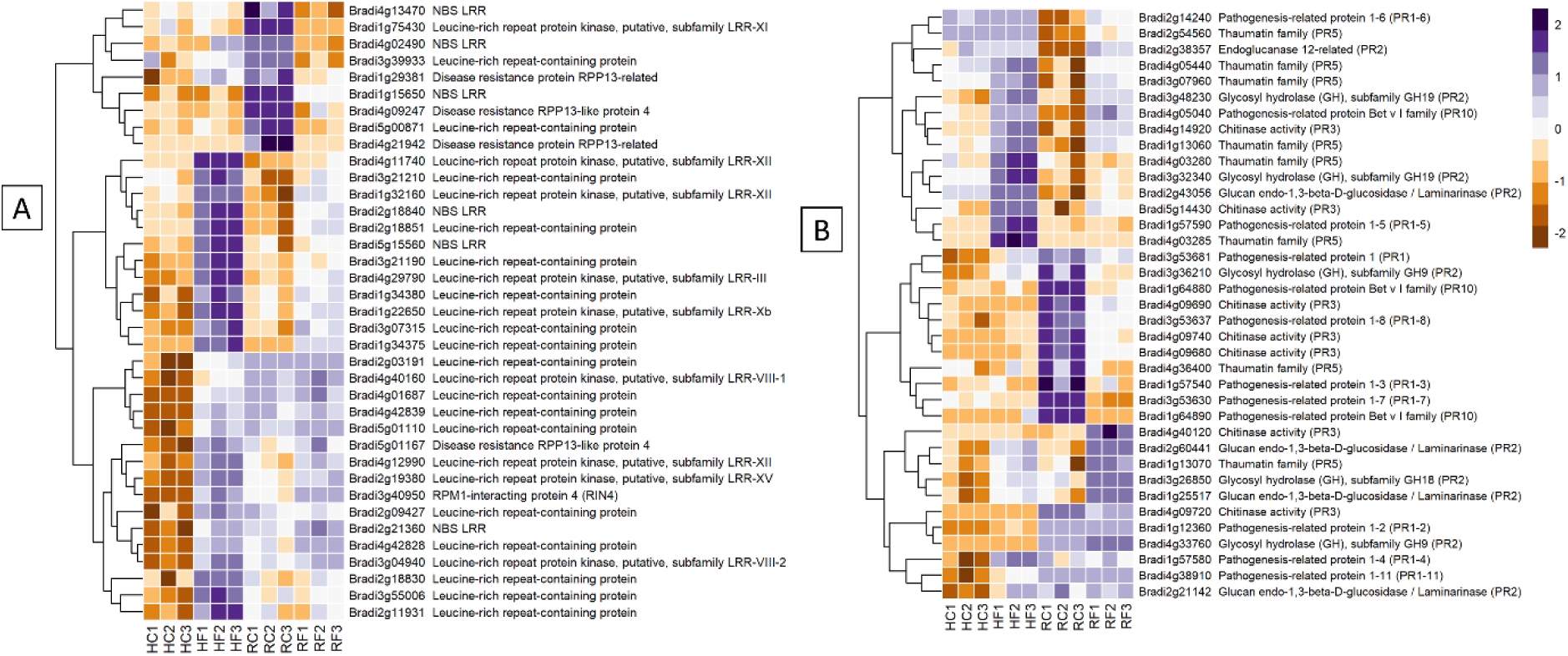
The most expressed or repressed receptor-related. (**A**) and defence-associated (**B**) *B. distachyon* genes. These genes showed a log-fold change of -3 ≤ Log2 ≥ 3 (**A**) and -2 ≤ Log2 ≥ 2 (**B**) with a p-adj < 0.05 in response to either FHB, FRR, or both. A more restrictive Log-fold threshold is used for (A) due to a larger number of significantly expressed or repressed receptor-related genes. The scale bar on the right is the Z-score for all the heatmaps. Three biological replicates for each of the four treatments are displayed as columns abbreviated as HC (Head-FHB control), HF (Head-FHB fungus), RC (Root-FRR Control), RF (Root-FRR fungus). The control samples (HC1-HC3, RC1-RC3) are normalised transcript counts from mock inoculated head (Water with Tween 20) and root (PDA slurry) *B. distachyon* tissues and were separately analysed through the RNA-seq pipeline with the respective inoculated sample tissues.

Increased expression of most of the WAK-related genes in FRR was only moderate (being below the Log Fold 2 threshold (Supplementary Table S7)).

### Genes encoding PRs and other antimicrobial compounds

Pathogenesis-related (*PR*) genes are important constituents of resistance because of their antimicrobial properties (Stintzi et al., 1993). *PR* genes were either similarly or differently expressed between FHB and FRR depending on the *PR* class (Figure 4B). Most of the *B. distachyon* PR1 genes described were differentially expressed between FHB and FRR (Figure 4B). The *PR* gene Bradi3g53681 was upregulated in FHB but downregulated in FRR. Most PR genes identified (Figure 4B) were also categorised based on hormone responsiveness, with majority being responsive to JA (Kouzai et al., 2016). Glucanases (*PR2*) (e.g. Bradi2g60441) and chitinases (*PR3*) (e.g. Bradi5g14430) on the other hand were generally expressed in both tissues. However, there was a cluster of chitinase genes on chromosome 4 that were significantly downregulated in FRR only (Figure 4B, Table S7). *PR5* genes encode thaumatin and most *PR5* genes were upregulated in FHB and/or FRR including Bradi1g13060 and Bradi1g13070. Bradi4g36400 was the only *PR5* gene downregulated in FRR (Supplementary Table S7). *PR10*/Bet v 1 gene products are also involved in pathogen resistance (Agarwal et al. 2016). One *PR10* gene (Bradi4g05040) was highly expressed in FHB and FRR while two *PR10* genes (Bradi1g64890 and Bradi1g64880) were exclusively downregulated response to FRR (Figure 4B).

Several other genes with predicted functions in secondary metabolite biosynthesis were identified (Supplementary Table S7). Those that were generally similarly differentially upregulated in both FHB and FRR include agmatine coumaroyltransferase (e.g. Bradi5g25166, Bradi3g23280, Bradi1g72220), phenylalanine ammonia lyase (e.g. Bradi3g47110, Bradi3g47120), tryptophan biosynthesis (Bradi1g55440, Bradi1g35600), and salt-stress/antifungal genes (Bradi1g25697, Bradi1g25552, Bradi5g03937) (Supplementary Table S7). There were, however, groups that showed different differential expression between FHB and FRR. These include flavonoid-related genes such as Bradi3g15700 and Bradi3g15690 that were highly downregulated in FRR. A cluster of six genes on chromosome 2 encoding Secologanin synthase-like genes were highly expressed in FHB but showed low or no expression in FRR (Supplementary Table S7). Lastly, one terpene related gene (Bradi3g35027) coding for alpha-humulene synthase was upregulated in FHB and downregulated in FRR (Supplementary Table S4).

### Genes encoding proteins with structural and cell wall-modification functions

Genes involved in cell-wall modification were one of the most differentially expressed groups between FHB and FRR. Of all the structural genes, expansins showed the most substantial difference between FHB and FRR (Figure 5A). The majority of identifiable expansins were highly downregulated in FRR and were generally not expressed in FHB (Figure 5A, Supplementary Table S7). The most downregulated of these include Bradi5g04120 and Bradi3g27460. The downregulation of expansin genes in response to FRR may, in part reflect the very high read count of these genes in non-inoculated root samples (Figure 5A, Supplementary Table S7). Xyloglucan-related genes were also differentially expressed between FHB and FRR, for example, Bradi3g31767 was upregulated in FHB and highly downregulated in FRR (Figure 5A, Supplementary Table S7). Cellulose synthase related genes Bradi3g00491 and Bradi4g33090 were expressed in FHB whereas two others, Bradi3g34490 and Bradi1g25130, were downregulated in FRR (Figure 5A, Supplementary Table S7). Most pectin-associated genes involved in de-esterification of pectin (pectinesterase, pectinmethylesterase, pectinmethylesterase inhibitor) were also differentially expressed. Aside from the pectin methylesterase genes Bradi2g11850 Bradi3g24750, the remaining functionally related genes were not significantly differentially expressed in FHB. In contrast, all pectin-associated genes were either upregulated or downregulated in FRR. The lignin biosynthesis-related gene, cinnamoyl-CoA reductase (Bradi3g19670), was upregulated in both FHB and FRR (Figure 5A).

**Figure 5:**
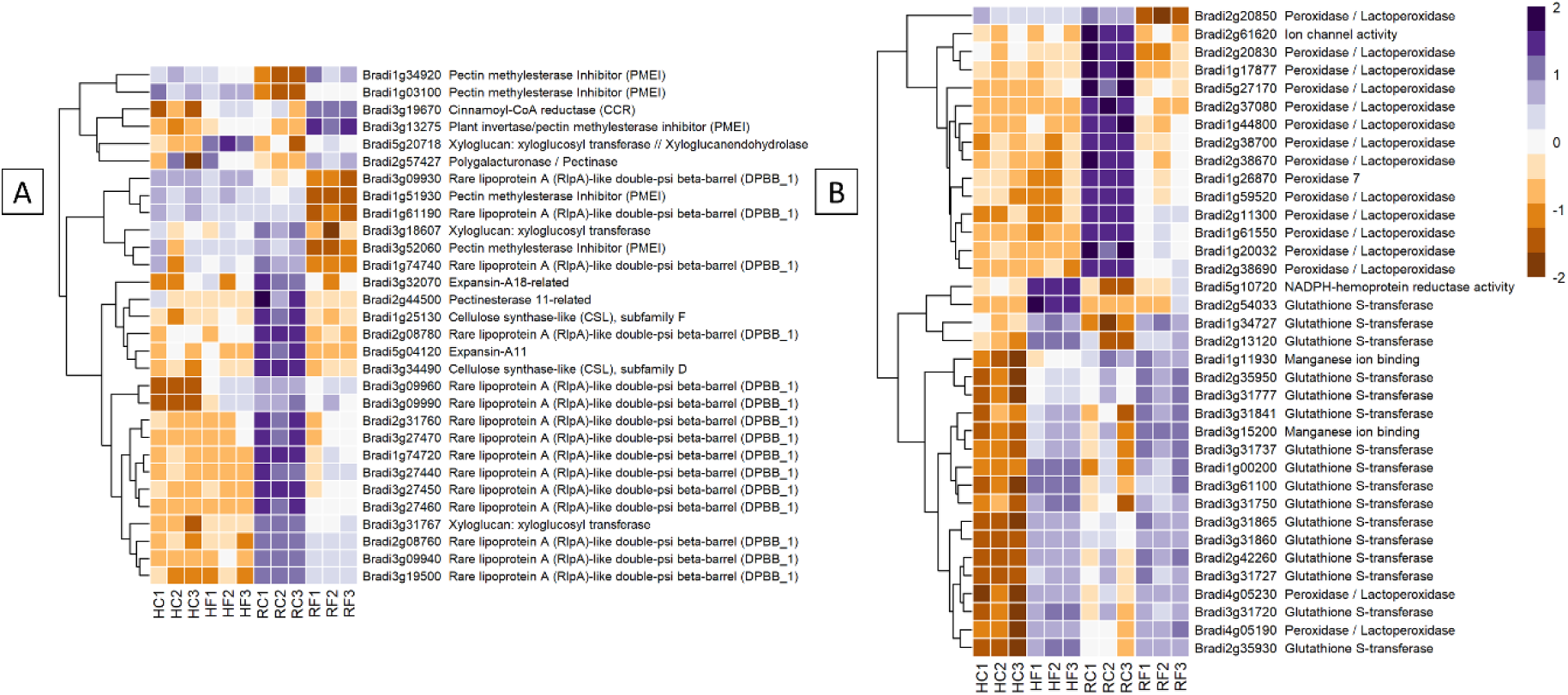
The most expressed or repressed cell-wall modification. (**A**) and ROS-associated (**B**) *B. distachyon* genes. These genes showed a log-fold change of -3 ≤ Log2 ≥ 3 (**A**) and -5 ≤ Log2 ≥ 5 (**B**) with a p-adj < 0.05 in response to either FHB, FRR, or both. A more restrictive Log-fold threshold is used for (A) due to a larger number of significantly expressed or repressed ROS-associated genes. The scale bar on the right is the Z-score for all the heatmaps. Three biological replicates for each of the four treatments are displayed as columns abbreviated as HC (Head-FHB control), HF (Head-FHB fungus), RC (Root-FRR Control), RF (Root-FRR fungus). The control samples (HC1-HC3, RC1-RC3) are normalised transcript counts from mock inoculated head (Water with Tween 20) and root (PDA slurry) *B. distachyon* tissues and were separately analysed through the RNA-seq pipeline with the respective inoculated sample tissues.

### Reactive oxygen species (ROS) - associated genes

Genes associated with reactive oxygen species (ROS) and antioxidation were the most differentially expressed group of genes between FHB and FRR (Figure 5B). The number of peroxidases genes downregulated during early FRR was the most for any gene class. Out of 68 genes, 8 peroxidases (e.g. Bradi3g10460, Bradi1g26870, and Bradi2g09650) and 33 peroxidases/lactoperoxidases (e.g. Bradi1g17877, Bradi1g59520, and Bradi2g38690) were exclusively downregulated in FRR (Figure 5B, Supp Table S7). No peroxidase genes were downregulated in FHB (Log Fold > 2) and some were upregulated (Supp Table S7). The difference in response of expression of peroxidases between FHB and FRR may be partly explained by the very high read count of peroxidase genes in non-inoculated root samples (Figure 5B, Table S1). Several glutathione-associated genes were upregulated in response to both FHB and FRR (Bradi1g34727, Bradi3g31841, Bradi2g35950, Supplementary Table S7). Most of the glutathione-associated genes, however, were exclusively upregulated in FHB being either downregulated or not significantly differentially expressed in FRR. Two classes of genes have been proposed as having roles in ROS metabolism of FCR (Powell et al. 2017). These include germin-like proteins (Bradi1g04907, Bradi3g15200, Bradi3g15190) and oxalate oxidase genes (Bradi1g11930, Bradi1g11920). These genes were highly upregulated during FHB but were not differentially expressed in FRR (Figure 5B, Supp Table S7). Likewise, expression of four NADPH-hemoprotein reductase activity genes was upregulated in FHB (Supplementary Table S7). Finally, a few *RBOHD* genes were also upregulated exclusively in FHB except for Bradi4g05540 which was upregulated exclusively in FRR.

### Transcription factors

Transcription factors showed different expression patterns between FHB and FRR tissues, depending on the class of transcription factor (Supplementary Table S7). The majority of *WRKY*, *NAC*, and *MADS* transcription factors were upregulated in FHB but not differentially expressed in FRR (Supplementary Table S7). The genes Bradi2g53760, Bradi2g00280, Bradi4g01950, and Bradi3g09810 encoding *WRKY* transcription factors were significantly expressed in FHB only. *MYB*, *AP2* domain, *EREBP*, and *ERF* transcription factors were broadly upregulated in both FHB and FRR (e.g. Bradi2g38560, Bradi3g18070, Bradi5g17490, Bradi3g12567), however many were not upregulated in FRR (Supplementary Table S7). The *bZIP* (e.g. Bradi2g50220, Bradi3g09340, Bradi2g06790) and *bHLH* transcription factors (e.g. Bradi2g12490, Bradi1g70860, and Bradi5g23580) were generally downregulated in FRR but most showed no large change in expression in FHB (Supplementary Table S7).

### Genes coding for ABC transporters and putative DON detoxification functional genes

DON detoxification is an important defence strategy to restrict Fusarium infection (Boutigny et al., 2008, Pasquet et al., 2016). The gene Bradi5g03300 encoding a UDP-glycosyltransferase can detoxify DON (Pasquet et al., 2016, Poppenberger et al., 2003). Several UDP glycosyltransferases, including Bradi5g03300, were highly expressed in FHB and but showed either no significant expression or moderate upregulation (Log Fold > 1) in FRR (Supplementary Table S7).

ATP-binding cassette (ABC) transporters have important roles in defence and have been found to be involved in resistance to FHB (Walter et al. 2015). Several ABC transporters were identified in *B. distachyon* that increase in expression in response to FCR (Powell et al., 2017b). Seven out of the nine identified were upregulated in FHB but were not expressed in FRR (e.g. Bradi3g35390, Bradi2g04577, Bradi5g03460) (Supplementary Table S7).

### Time-course analysis of differentially expressed *Brachypodium* genes

The FHB and FRR assays used to generate the RNA-seq data were repeated with the addition of two additional time-points and expression of a selection of genes were assessed using RT-qPCR to confirm the results from RNA-seq and to provide additional information on temporal aspects of expression. The selected genes were all differentially expressed between FHB and FRR in the RNA-seq except for *WRKY45* (Bradi2g30695) which was not expressed in either FHB or FRR. These genes either have predicted roles with phytohormones or have roles in plant defence. In almost every instance, the expression differential at the first time point (Figure 6), was similar to that observed in the RNA-seq experiment (Supplementary Table S4). The results from the RT-qPCR confirmed those from the RNA-seq and showed that, for many of these genes, the expression differences between FHB and FRR were maintained over time. The SA-responsive *WRKY45* (Bradi2g30695), SA-associated *MES1* (Bradi2g52110), auxin responsive homolog *OsAUX/IAA* (Bradi1g09090), cytokinin-responsive homolog *OsLOG1* (Bradi2g42190), cytokinin-responsive homolog *OsA-RR9* (Bradi4g43090), and defence-associated *ABC* transporter (Bradi2g43120) showed differential expression patterns between FHB and FRR over the entire time course (Figure 6). In contrast, the genes JA and SA-responsive *PR1-5* (Bradi1g57590), SA-associated *NPR4* (Bradi2g54340), auxin-responsive homolog of *OsGH3* (Bradi2g50840), the orthologue of *TaCYP450* with a role in DON detoxification (Bradi2g44150), and defence-associated oxalate oxidase (Bradi1g11930) showed similar expression patterns at 3 dpi and 5 dpi for FHB and FRR (Figure 6) despite being differentially expressed at the earlier time points in the RNA-seq data (Table S4). The JA-associated genes *LOX2* (Bradi3g39980), and *JAZ* (Bradi4g31240) which were similarly expressed in FHB and FRR in the RNA-seq (Supplementary Table S6) were also similarly expressed at 3 dpi (Supplementary Figure S1).

**Figure 6:**
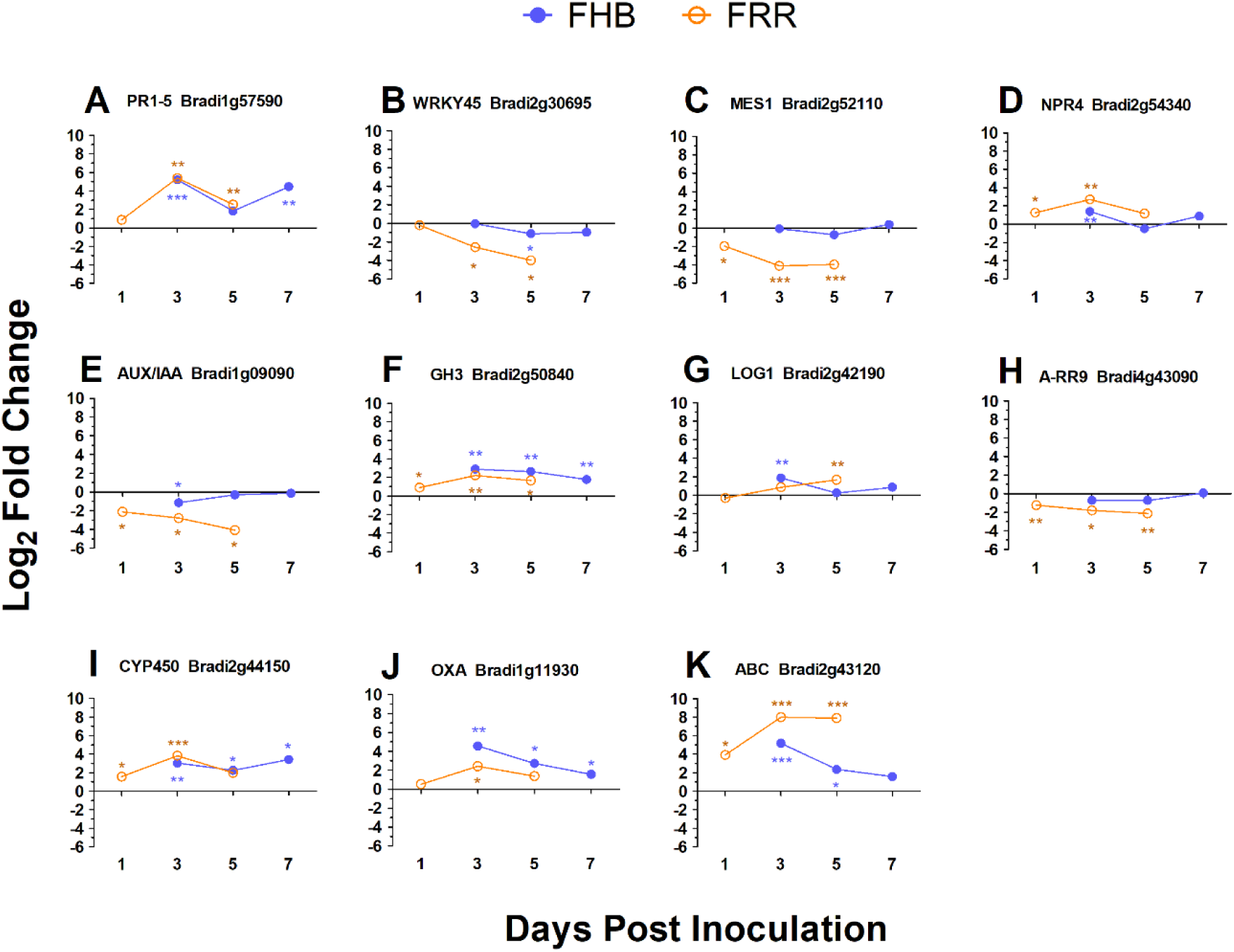
Expression time-course of differentially expressed *B. distachyon* genes in FHB and FRR. The blue lines denote FHB and orange lines denote FRR. The Log values presented are calculated by comparing infected tissue against mock-inoculated treatments. The reference housekeeping gene *GAPDH* was used. Each point is the average of three biological replicates and 2-3 technical replicates. Levels of significance relative to the mock-control, Cq t-test *p<0.05, **p<0.01, *** p < 0.001.

### Differential expression of *F. graminearum* genes and predicted effectors

Gene transcript counts of *F. graminearum* in the FHB and FRR samples were compared against *F. graminearum in vitro* control samples to determine whether gene expression in the pathogen differed when infecting the two tissues (Tables S8 and S9). Differentially expressed genes were compared between FHB and FRR (Supplementary Table S10). Moderate proportions of the transcript reads mapped to the *F. graminearum* PH1 assembly, with 8% to 15% for FHB and FRR respectively. A total of 4,567 *F. graminearum* genes were significantly responsive in FHB or FRR, or in both (Figure 7A), however only 6% of these were functionally characterised on UniProt (Consortium, 2018). A total of 3,499 genes were significantly differentially expressed in spike infection (FHB), of which 1,919 (55%) were upregulated (Figure 7A), while 3,214 genes were differentially expressed during root infection (FRR), of which 1,815 (56%) were upregulated (Figure 7A). From the significantly differentially expressed *F. graminearum* genes, 1,167 (26%) were upregulated and 934 (21%) were downregulated in both root and spike tissues, respectively, relative to axenic culture medium (Figure 7A). Only 45 (1%) genes were upregulated in one tissue but downregulated in the other (Figure 7A). Gene-list enrichment was performed on all significantly upregulated and downregulated genes (Supplementary Table S2). Most pathways were similarly expressed between FHB and FRR (e.g. ‘metabolic pathways’, ‘pentose and glucuronate interconversion’, and ‘starch and sucrose metabolism’). The pathways ‘cyanoamino acid metabolism’ and ‘other glycan degradation’ were, however, exclusively upregulated in FHB whereas ‘arginine and proline metabolism’ was exclusively upregulated in FRR. No pathways were significantly downregulated in either FHB or FRR.

**Figure 7:**
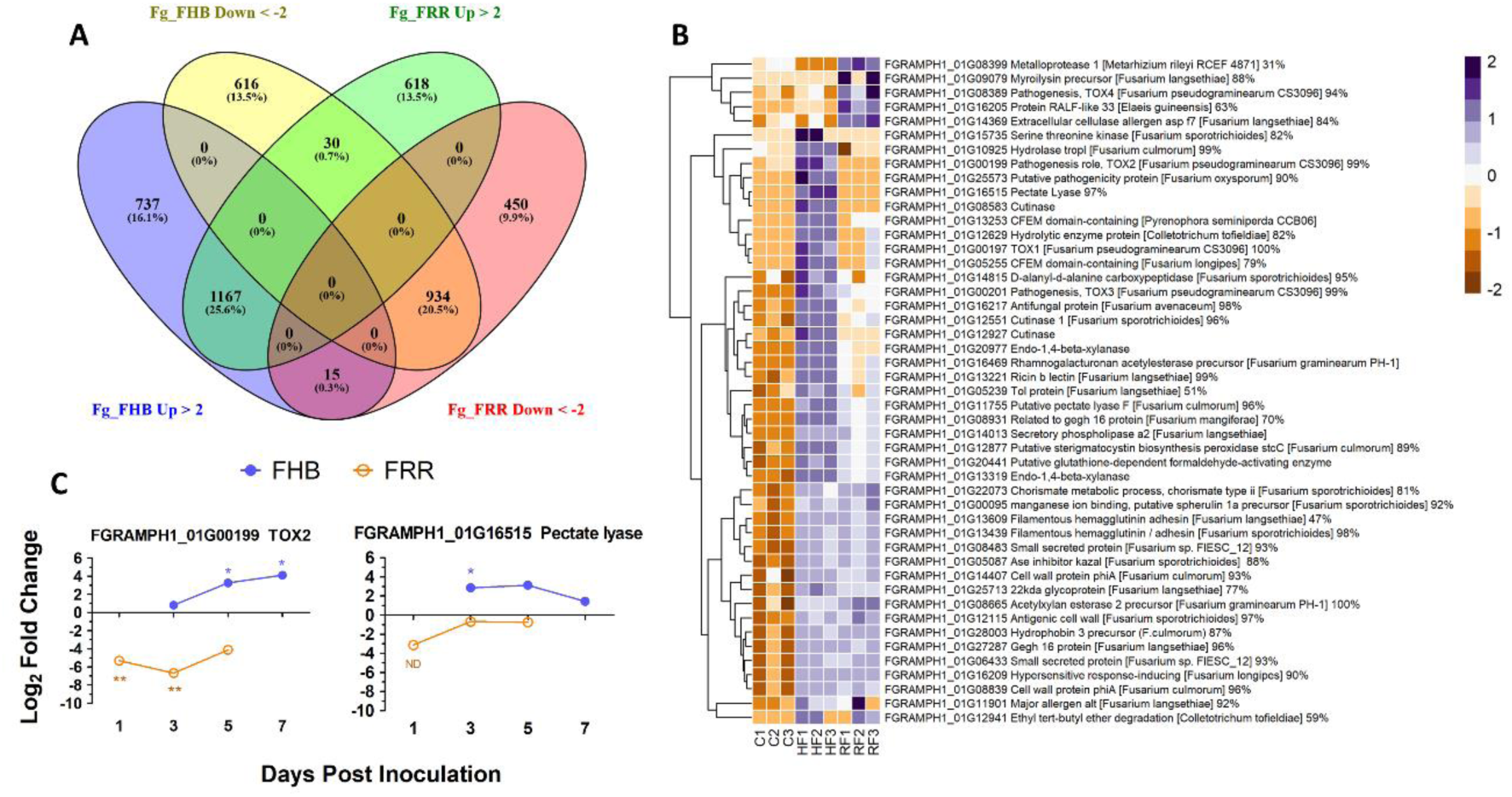
The differential expression of F. graminearum genes between FHB and FRR. (**A**). A summary of all significantly upregulated or downregulated F. graminearum genes in response to FHB and FRR. The threshold of -2 ≤ x ≥ 2 Log-fold change and p-adj < 0.05 was applied to all genes. (**B**) The most upregulated F. graminearum predicted-effector genes in FHB and FRR that have predicted functions. These genes have an expression of Log-fold increase ≥ 3 in FHB and/or FRR. Three biological replicates for each of the three treatments are displayed as columns. Control Treatment (C1-3) is the average normalised gene counts (two values per column/biological replicate) of in vitro treatments (four-day-old samples grown in Czapek-Dox Liquid media). The same in vitro sample was analysed separately (DESeq) with the FHB and FRR samples. (**C**) Expression time-course of two differentially expressed F. graminearum genes (FGRAMPH1_01G00199 (TOX2) and FGRAMPH1_01G16515 (pectate lyase)) in FHB and FRR identified in **B**. Blue lines with solid circles denote FHB and orange lines with open circles denote FRR. Log values presented are calculated by comparing infected tissue against mock-inoculated treatments. The housekeeping gene GzUBH was used. Level of significance relative to the mock-control: Cq t-test *p < 0.05, **p < 0.01. “ND” denotes no statistical comparison due to insufficient data points in treatment. Abbreviations: Bd (B. distachyon), Up (Upregulated), Down (Downregulated), FRR (Fusarium Root Rot), FHB (Fusarium Head Blight). C (in vitro control respective to FHB and FRR), HF (Head-FHB fungus), RF (Root-FRR fungus).

All the significantly expressed genes were filtered by association with the *F. graminearum* secretome database (Brown et al., 2012). Only 92 (3%) of FHB-responsive genes (Figure 7A) were classed as effectors and 37 (40%) of these were exclusively expressed or repressed in FHB. On the other hand, 70 (2%) of FRR-responsive genes (Figure 7A) were classed as effectors and 15 (21%) of these were exclusively expressed or repressed in root tissues (FRR) (Supplementary Table S11). Fifty-eight (54%) of all these effector-associated genes had a predicted function (From *F. graminearum* (UniProt (Consortium, 2018)) or through protein homology in different fungal species (BLAST (Sayers et al., 2020), Supplementary Table S11). The upregulated genes were then selected since they were predicted to be the effectors playing important roles in FHB and FRR virulence. A total of 80 *F. graminearum* genes were highly upregulated (Log2-fold ≥ 3) in FHB and/or FRR (Table S11) and 47 of these with predicted functions are presented (Figure 7B). Many of the predicted functions were associated with cell-wall degradation and pathogenesis. The majority of *F. graminearum* predicted effectors were similarly expressed between FHB and FRR (Figure 7B). The predicted effectors FGRAMPH1_01G14013 (secretory phospholipase), FGRAMPH1_01G20977 (Endo- 1,4-beta-xylanase), FGRAMPH1_01G16469 (rhamnogalacturonan acetylesterase precursor), FGRAMPH1_01G13253 (CFEM domain-containing), and FGRAMPH1_01G27287 (gEgh 16 protein) were among the most highly upregulated effectors in both FHB and FRR. A number of effectors, however, were upregulated in a tissue-specific manner. Nine predicted effectors (e.g. FGRAMPH1_01G00199 (*TOX2*), FGRAMPH1_01G16515 (Pectate Lyase), and FGRAMPH1_01G12927 (Cutinase)) were exclusively upregulated in FHB, whereas 5 (e.g. FGRAMPH1_01G08389 (*TOX4*), FGRAMPH1_01G08399 (Metalloprotease), and FGRAMPH1_01G09079 (myroilysin precursor)) were exclusively upregulated in FRR (Figure 7B).

The expression of two effector genes (FGRAMPH1_01G00199 (*TOX2*) and FGRAMPH1_01G16515 (Pectate Lyase)) that showed differential expression between FHB and FRR in the RNA-seq data set (Supplementary Table S11) were examined using RT-qPCR (Figure 7C). Although the absolute expression values differed between experiments for the same gene at 1 dpi for FRR and 3 dpi for FHB (Figure 7B and 6C), the differential expression between tissues was maintained over time (Figure 7C). *TOX2* was downregulated in FRR but upregulated in FHB whereas pectate lyase was only significantly upregulated in FHB at 3 dpi (Figure 7C).

Trichothecene production is regulated by the *Tri* gene cluster (Kimura et al., 2007, Kimura et al., 2003). *F. graminearum* PH1 is a 15-acetylDON (ADON) producer (Kimura et al., 2007).

The expression of the *Tri5*-gene cluster was investigated to identify any difference in transcription of DON associated genes between FHB and FRR. The essential DON biosynthetic triplet of genes (*Tri4*, *Tri5*, *Tri11*) were the most upregulated genes in both FHB and FRR (Table. 2). The transcriptional regulators (*Tri6*, *Tri10*), and *Tri12* transporter were also upregulated in both FHB and FRR and there were also high levels of expression of *Tri3*, *Tri9* and *Tri14* in both tissues (Table 1). Despite the low but significant expression in FHB, *Tri8* was the only gene in the Tri5-gene cluster that was exclusively expressed in FHB (Table 1). *Tri8* encodes a C-3 deacetylase that is involved in 15-ADON production (Alexander et al., 2011). The *Tri7* and *Tri13* genes in *F. graminearum* PH1 do not encode functional proteins as this isolate is a 15-AcDON chemotype and these genes are involved in the synthesis of nivalenol (Kimura et al., 2003, Lee et al., 2002). Neither of these genes exceeded the significance log-fold threshold in either FHB or FRR (Table 1).

**Table 1:**
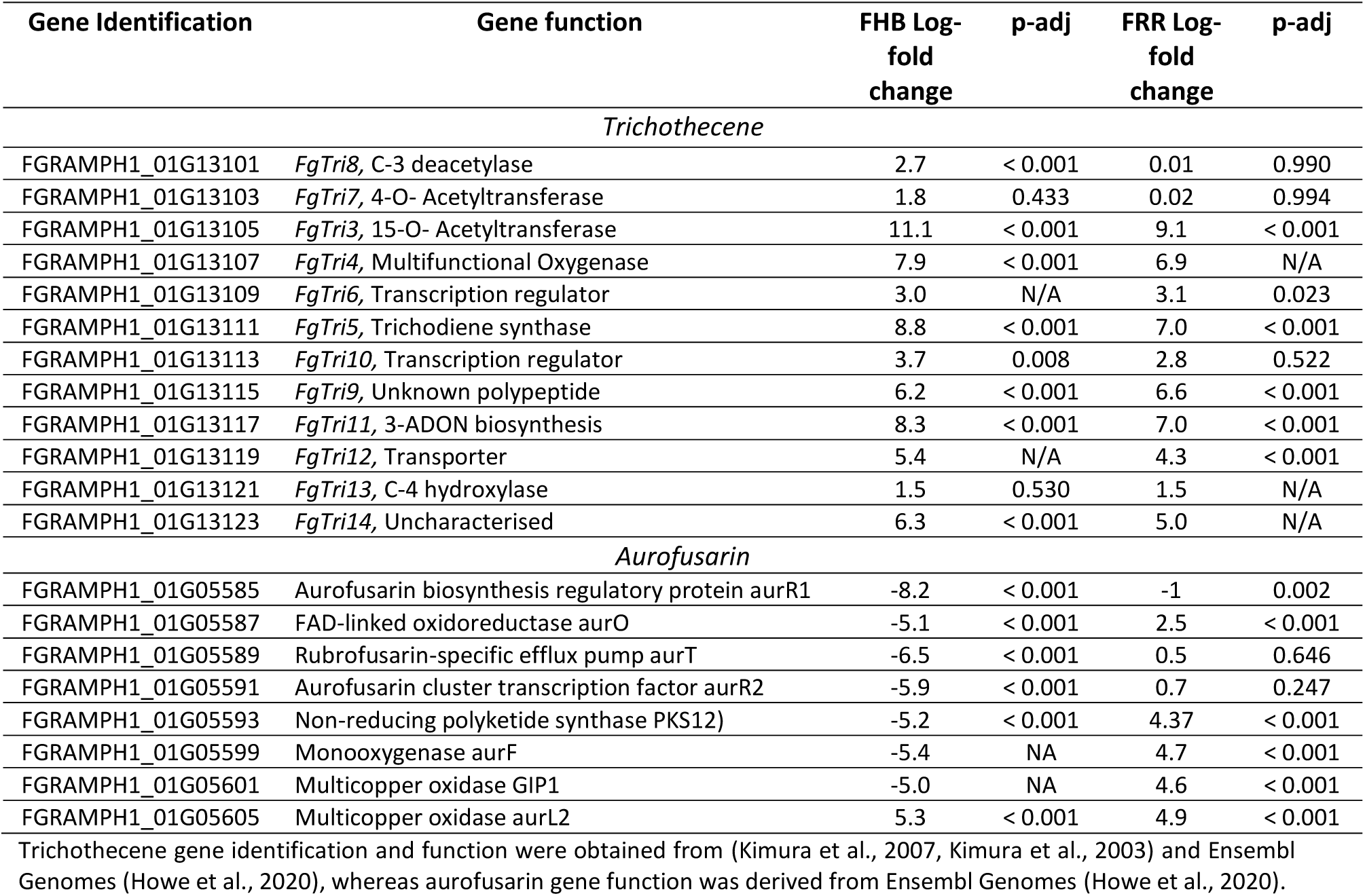
Notable *F. graminearum* secondary metabolite clusters that were significantly expressed in FHB and FRR.

The cluster of genes involved in the synthesis of the red pigment aurofusarin were differentially expressed between FHB and FRR (Table 1). Most of the genes in the pathway were significantly downregulated in spike tissue (FHB) but upregulated in root tissue (FRR) (Table 1). Both FGRAMPH1_01G05593 (*PKS12*) and FGRAMPH1_01G05587 (*aurO*) (Table 1) were among the 30 genes that were highly upregulated in FRR and downregulated in FHB (Figure 7A). FGRAMPH1_01G05599 (*aurF*) and FGRAMPH1_01G05601 (*GIP1*) also displayed differential in expression but the effect was not statistically significant for FHB (Table 1). The exceptions to this differential trend were FGRAMPH1_01G05605 (*aurL2*) which was significantly upregulated in both FHB and FRR, and FGRAMPH1_01G05589 (*aurT*) and FGRAMPH1_01G05591 (*aurR2*) which were downregulated in response to FHB but were not significantly upregulated in response to FRR (Table 1). Likewise, FGRAMPH1_01G05585 (*aurR1*) was significantly downregulated relative to *in vitro* conditions in both FHB and FRR, but to a lesser degree in FRR (Table 1).

## Discussion

The transcriptome of *B. distachyon* exhibited many tissue-specific differences in response to infection of spike and root tissues by *F. graminearum*. The majority of differentially expressed genes were exclusively expressed or repressed in response to FHB or FRR (Figure 1C). Interestingly, while an equal number of genes were upregulated or downregulated in response to FRR (Figure 1A) the vast majority of the differentially expressed genes in FHB were upregulated (Figure 1A). The preponderance of upregulation over suppression of gene expression in response to FHB has also been reported with wheat (Buerstmayr et al., 2021). More genes were upregulated than downregulated in *B. distachyon* following infection of the base of seedlings (Fusarium Crown Rot (FCR)) by *F. pseudograminearum* (Powell et al., 2017a, Powell et al., 2017b) suggesting that the transcriptional response of FCR may be more like that to FHB than to FRR. Differences in the proportion of up and downregulated genes in different tissues has been observed in other host-pathogen interactions. Expression of more genes in *A. thaliana* roots was repressed in response to *F. oxysporum* infection than in response to leaf infection (Chen et al., 2014) and approximately half of the differentially expressed genes displayed root or leaf-specific expression following infection of *A. thaliana* by *F. oxysporum* (Lyons et al., 2015). Many pathogens exhibit organ specificity being able to infect some tissues but not others but it is unclear whether these effects are controlled by the host or pathogen (Strugala et al., 2015). The rice cultivar Tai-Nong is resistant to foliar infection by isolate 031 of *Magnaporthe oryzae* but is susceptible to root infection by the same isolate (Jansen et al., 2006). Additionally, root systems of the FHB-resistant wheat cv. Sumai 3 were found to be susceptible to FRR (Wang et al., 2015). In the present study we provide evidence of tissue-specific host responses to infection by *F. graminearum* as well as tissue-specific gene expression in the pathogen.

We previously showed that exogenous application of phytohormones induced tissue-specific effects on resistance to FHB and FRR in *B. distachyon* (Haidoulis and Nicholson, 2020) indicating that pathways controlled by phytohormones play different roles in resistance in spike and root tissues. More JA-responsive genes were significantly expressed and to a greater extent in FHB than in FRR (Figure 2, Figure 3A, Supplementary Table S6), although there were similar changes in expression of some biosynthesis and signalling genes (*LOX* and *JAZ*) in FHB and FRR (Supplementary Table S6, Supplementary Figure S1). For most biosynthesis genes (*OPR* and *LOX*), the basal level of expression was much higher in non- inoculated root tissues compared to non-inoculated spikes (Supplementary Table S6) which may account for the apparent reduced level of expression of JA-associated processes during infection in roots. Supporting these results, *LOX* and *JAZ* genes were also found to be upregulated in *B. distachyon* in response to FHB, FCR, or FRR (Pasquet et al., 2014, Powell et al., 2017b, Ding et al., 2021), in wheat to FHB (Sun et al., 2016, Pan et al., 2018, Wang et al., 2018b), and in wheat to FCR and FRR (Wang et al., 2018c, Powell et al., 2017a). Similar to JA, ethylene biosynthesis (*AOC*) and many signalling genes (*ERF1* and AP2 domain containing genes) were upregulated in both FHB and FRR (Figure 3B, Supp Table S6). Genes with the same function were also upregulated in *B. distachyon* FCR and FRR (Powell et al., 2017b, Ding et al., 2021), and wheat FHB (Sun et al., 2016, Pasquet et al., 2014). Ethylene functions synergistically with JA signalling in plants by prioritizing JA signalling and fine-tuning resistance to necrotrophic pathogens (Bari and Jones, 2009, Pieterse et al., 2012) via *ERF1/ORA59* transcription factors (Lorenzo et al., 2003, Pieterse et al., 2012, McGrath et al., 2005). Given the abundance of ethylene-related transcription factors upregulated in both FHB and FRR (Figure 3B) and the overrepresentation of ethylene-associated processes in FHB and FRR (Figure 2), the data suggests that the ethylene branch of JA signalling is activated in both tissues and that the JA/ethylene synergism is important in FHB and FRR. Supporting this, exogenous application of JA and the ethylene precursor 1-aminocyclopropane-1-carboxylic acid (ACC) had similar effects on resistance in both tissues (Haidoulis and Nicholson, 2020). In the present study, JA and ethylene - related gene expression was broadly similar in FHB and FRR (Figure 3A and Figure 3B). There is a contrast in the effects of exogenous phytohormone application and gene expression in the different tissues for the same hormones (Supplementary Table S12). This may be caused by the different native states of expression of the phytohormone associated genes in spikes and roots which may dictate whether associated genes are significantly expressed or not on exposure to the pathogen. Alternatively, this may be due to a potentially contrasting lifestyle of *F. graminearum* in FHB and FRR. We previously suggested that *F. graminearum* may be behaving predominantly as a necrotroph in *B. distachyon* roots (Haidoulis and Nicholson, 2020). This may trigger a different host response in *B. distachyon* roots.

In contrast to JA and ethylene, there was minimal evidence for altered expression of genes involved in SA biosynthesis or signalling in FHB and FRR (Supplementary Table S6). The key SA-negative regulator *NPR4* (Figure 6D) (Ding et al., 2018), and the SA-responsive *WRKY* transcription factor (Figure 6B) (Kakei et al., 2015, Kouzai et al., 2016) were upregulated and downregulated, respectively, in FHB and FRR. Together, these findings indicate that the JA, rather than the SA, pathway is predominantly expressed in both tissues at the onset of first symptoms. An absence or a very low amount of SA-related gene transcription was also identified in wheat, *B. distachyon*, and *Arabidopsis* in response to *Fusarium* infection (Lyons et al., 2015, Sun et al., 2016, Powell et al., 2017b). However, in contrast, SA signalling was suggested to be a significant factor in the *B. distachyon* response to FRR (Ding et al., 2021). The difference may be due to FRR sampling which was 4 dpi, later than in this study.

Unlike the canonical defence-associated phytohormones, expression of auxin- and cytokinin-associated genes generally differed between FHB and FRR. For auxin, two GO processes involved in indole-containing compound biosynthesis and metabolic processes were overrepresented in FHB but not in FRR (Figure 2). The signalling genes encoding *AUX/IAA*, *ARF*, and *SAUR* were generally upregulated in response to FHB but suppressed in FRR (Figure 3C, Figure 6E). Two of the same *ARF*s and a *SAUR* gene were also reported to be upregulated in FHB in *B. distachyon* (Pasquet et al., 2014), while two SAUR genes were similarly downregulated in FRR of *B. distachyon* (Ding et al., 2021). In contrast, auxin homeostasis-associated gene *GH3* genes (Staswick et al., 2005) showed similar expression at 3 dpi and 5 dpi in FHB and FRR (Figure 6F). Rice *GH3* homologues are associated with resistance responses (Domingo et al., 2009, Fu et al., 2011) and biosynthesis and auxin-related signalling can affect resistance to plant pathogens (Bari and Jones, 2009, Kazan and Manners, 2009). Lyons and colleagues (2015) concluded that auxins were important components of defence responses to *F. oxysporum* in both shoot and root tissues of Arabidopsis. We reported previously that exogenous application of auxin increased resistance to both FHB and FRR in *distachyon* (Haidoulis and Nicholson, 2020) and others have found similar effects on FHB in barley (Petti et al., 2012). In contrast, there is evidence that *F. graminearum*-induced FHB and leaf susceptibility increased following exogenous IAA application in wheat (Su et al., 2020). Furthermore, auxin content and signalling have been associated with FHB susceptibility (Wang et al., 2018a, Brauer et al., 2019). Disruption of the auxin receptor gene *TaTIR1* and attenuation of *TaARF2* expression in wheat reduced susceptibility to FHB (Su et al., 2020, Chen et al., 2016). Auxin homeostasis may be important in resistance to FHB and FRR and exogenous application of auxin or disruption of signalling may both suppress the potential of the pathogen to manipulate levels of auxin in the host. This possibility is supported by the finding that *F. graminearum* can synthesise auxin (Luo et al., 2016).

Aside from a study by (Powell et al., 2017a), the role of cytokinins in *F. graminearum* infection has not been investigated beyond the exogenous application of cytokinins which enhanced FHB and FRR susceptibility (Haidoulis and Nicholson, 2020). Most biosynthesis genes (*LOG1*, *IPT*, *UGT85A1*) and signalling genes (*Type A RR*s and *B RR*s) were differentially expressed between the two tissues in response to *F. graminearum* (Figure 3D, Figure 6G, and Figure 6H), highlighting a difference in response to FHB and FRR in terms of cytokinin signalling. Both auxin and cytokinin-associated signalling genes tended to be upregulated in FHB but downregulated in FRR although exogenous application of these phytohormones had similar but contrasting effects on resistance in both tissues with auxin increasing resistance while cytokinin reducing resistance to both FHB and FRR (Supplementary Table S12). This may reflect differences in the infection strategy of *F. graminearum* between tissues, discussed in (Haidoulis and Nicholson, 2020), however further research is needed to understand the role of auxin and cytokinin in the defence response to *F. graminearum*.

Several types of antimicrobial-associated genes were similarly expressed in FHB and FRR (Figure 4B). Five classes of *PR* genes were generally upregulated in both FHB and FRR (Figure 4B). Many of the same *PR* gene classes were similarly expressed in FHB of wheat (Pan et al., 2018, Pritsch et al., 2000, Wang et al., 2018a), FCR of wheat (Powell et al., 2017a), FHB of barley (Boddu et al., 2006), and FHB, FCR, and FRR of *B. distachyon* (Pasquet et al., 2014, Powell et al., 2017a, Ding et al., 2021). Together the data suggests that *PR* gene expression is a common host response to most *Fusarium* diseases.

ROS-associated genes and pathways were the most differently expressed in the dataset. Most peroxidases, involved in the breakdown of hydrogen peroxide, were downregulated in FRR (Supplementary Table S7, Figure 5B). Several of the same peroxidase genes were also highly downregulated in another study that investigated the response of *B. distachyon* to FRR (Ding et al., 2021). The overrepresented GO biological pathways comprising hydrogen peroxide catabolic process, ROS metabolic process, and response to oxidative stress. These processes were highly downregulated in FRR (Figure 2). The cellular oxidant detoxification process was exclusively downregulated in roots (Figure 2). Together this suggests that in FRR there is predominantly a reduction in ROS metabolism, oxidative response, and oxidant detoxification. This observation may, in part, be due to the high basal expression of peroxidases in non-inoculated roots samples compared to non-inoculated spikes (Figure 2). In contrast to FRR, most peroxidases were not significantly expressed in FHB (Supplementary Table S7, Figure 5B). However, genes involved in glutathione metabolism, mostly glutathione-S-transferases (GST) (Supplementary Table S5 and S7), were generally highly upregulated in FHB but only moderately upregulated in FRR (Figure 5B). Likewise, the GO process glutathione metabolic process was only overrepresented in FHB (Figure 2). Glutathione metabolism, was reported to be an important response to *B. distachyon* FCR (Powell et al., 2017b). Likewise several glutathione associated genes were upregulated in *B. distachyon* to FRR (Ding et al., 2021), and GST expression was upregulated in barley FHB (Boddu et al., 2006) and associated with resistance in wheat FHB (Pan et al., 2018). The high upregulation in FHB may be because of relatively low basal levels of GST in non-inoculated spikes compared to non-inoculated roots (Figure 2). There is, however, a notable difference in expression of these two different classes of enzymes with similar function between FHB and FRR (peroxidase and glutathione) which suggests a different ROS state between FHB and FRR.

It was possible to examine the transcriptomes of both *F. graminearum* and *B. distachyon* in the same material. This offered the opportunity to observe both the host and pathogen components of the interaction in the two tissues. Cell-wall degrading enzymes (CWDEs) are associated with both necrotrophic and hemibiotrophic pathogens (Zhao et al., 2014, Kabbage et al., 2015). *F. graminearum* is known to express an abundance of CWDE genes (Cuomo et al., 2007, Kikot et al., 2009) in symptomatic tissue (Brown et al., 2017). The role of CWDEs is likely as a means for nutrient acquisition and/or as effectors (Cuomo et al., 2007, Walton, 1994). Several CWDEs were among the putative *F. graminearum* effectors and increased in expression in both FHB and FRR (Figure 7B). Enhanced expression of CWDEs, including genes involved in pectin degradation (Figure 7B), have also been reported in several other studies with wheat, barley, and maize FHB (Lysøe et al., 2011, Harris et al., 2016, Brown et al., 2017, Pan et al., 2018) and *B. distachyon* FRR (Ding et al., 2022). An effective plant strategy to detect necrotrophic pathogens is the release of DAMPs like oligogalacturonides by wall-associated receptor kinases (WAKs) that bind galacturonan (Wang et al., 2014, He et al., 1998). WAKS serve as pectin debris receptors (Kohorn and Kohorn, 2012). Interestingly, expression of WAKs was generally only upregulated in response to FHB (Supplementary Table S7) despite similar expression of pectin-associated CDWEs by *F. graminearum* in both tissues (Figure 7B). Gadaleta and colleagues (2019) found a *TaWAK2* gene associated with the FHB resistance QTL QFhb.mgb-2A in wheat (Gadaleta et al., 2019). Elevated expression of several WAK genes was reported in Bd roots at 5 dpi (Ding et al., 2021) suggesting that there may be a delayed response in root tissues.

The core *Tri* genes for DON production (Kimura et al., 2003) were upregulated in both FHB and FRR (Table 1). This cluster has been shown to be upregulated in FHB in several species (Lysøe et al., 2011, Harris et al., 2016, Brown et al., 2017, Pan et al., 2018) and in *B. distachyon* FRR (Ding et al., 2022). Wang and colleagues also reported the presence of DON in root tissues (Wang et al., 2015). Although *F. graminearum* has been shown to produce DON in both *B. distachyon* floral and root tissue during infection, there is evidence that it does not act as a virulence factor in roots (Ding et al., 2022). Furthermore DON does not act as a virulence factor in FHB of barley and maize (Maier et al., 2006). It is plausible that the influence of DON on virulence is tissue-specific despite its tissue-independent production. DON detoxification during early infection is an effective host strategy to increase resistance against FHB and FCR (Mandalà et al., 2019). Several UDP-glycosyltransferase (UGT) genes were found to be upregulated in FHB in the present study (Supplementary Table S7) and upregulation of many of these was also observed in other studies with FHB, FCR, and FRR (Wang et al., 2018c, Powell et al., 2017b, Pasquet et al., 2014). UDP-glucosyltransferases can glucosylate DON into the less toxic deoxynivalenol-3-*O*-glucose form (Pasquet et al., 2016). Interestingly, the gene encoding a UDP-glucosyltransferase that was highly expressed in FHB (Bradi5g03300) (Supplementary Table S7) has been shown to confer spike resistance to *F. graminearum* and root tolerance to DON in *B. distachyon* (Pasquet et al., 2016) and type 2 FHB resistance when expressed in wheat (Gatti et al., 2019). Bradi2g44150 (Figure 6I) is a homolog of *TaCYP72A* involved in resistance to DON (Gunupuru et al., 2018), and was significantly upregulated in FHB in the RNA-seq and at 3 dpi in both tissues (Figure 6I, Supplementary Table S7). If this homolog serves a similar DON resistance function in *B. distachyon* as in wheat (Gunupuru et al., 2018), then activation of DON-detoxification processes may be occurring similarly in both tissues.

*F. graminearum* can behave as a facultative hemibiotroph in spike tissues of wheat (Brown et al., 2010). Effectors were originally believed to be biotroph-specific however evidence is accumulating to suggest their importance for hemibiotrophic and necrotrophic pathogens (Amselem et al., 2011, Guyon et al., 2014, Kabbage et al., 2015). Plants often detect effectors using NBS-NLRs or LRR-RLK receptors. The receptor signalling GO processes were overrepresented only in FHB (Figure 2). Furthermore, the differential expression of many NBS-LRRs and LRR-RLKs was observed to be tissue-dependent in the present study (Figure 4A). For example, homologues of the *Recognition of Peronospora 13* (RPP13) genes were differentially expressed between FHB and FRR (Figure 4A, Supplementary Table S7). *RPP13* confers resistance to diseases such as downy mildew (Bittner-Eddy et al., 2000), and homologues of this gene were reported to be upregulated in response to *B. distachyon* FCR (Powell et al., 2017b). It is unclear why there is such a marked difference in the expression of genes involved in pathogen recognition in the two tissues, but it suggests that the host may be responding to different sets of effectors produced by *F. graminearum* in root and spike tissues.

A comparison of the different *F. graminearum* genes and effector-like genes in FHB and FRR revealed evidence for tissue-specific gene expression of components of the secretome. Of the *F. graminearum* predicted effector genes, 42% and 24% were exclusively expressed in FHB and FRR, respectively (Supplementary Table S11). This suggests that the expression of a large proportion of the *F. graminearum* secretome genes are controlled in a tissue-specific manner following infection of *B. distachyon* tissues. Genes encoding cutinases are present in relatively large numbers in hemibiotrophic and necrotrophic pathogens such as *Gaeumannomyces graminis* and *Magnaporthe oryzae* (Zhao et al., 2014). Two of the three expressed *F. graminearum* cutinases were exclusively upregulated in FHB (Supplementary Table S11). This finding is supported by reports that cutinases were expressed in wheat and barley FHB (Lysøe et al., 2011, Harris et al., 2016, Pan et al., 2018), but not in *B. distachyon* FRR (Ding et al., 2022). This difference between tissues is likely due to the cutin layer being present only on the epidermis of shoot tissues (Walton, 1994). The expression of one putative cutinase in roots (FGRAMPH1_01G12551) may also be associated with a different pathogenicity-associated role (Walton, 1994). *Fusarium graminearum* can produce the naphthoquinone aurofusarin, which is a red pigment and is synthesised by the aurofusarin gene cluster (Malz et al., 2005, Frandsen et al., 2006). The cluster of genes involved in aurofusarin biosynthesis (excluding FGRAMPH1_01G05605 (FGSG_02330)) were downregulated in FHB but the key genes were upregulated in FRR (Table 1). A similar result was observed for symptomatic wheat FHB with the same *F. graminearum* isolate (Brown et al., 2017). This suggests aurofusarin is being produced by *F. graminearum* PH1 during FRR pathogenesis of *B. distachyon*. A potential role of aurofusarin in FRR is not known although it was shown to not affect FHB virulence in wheat and barley (Kim et al., 2005, Malz et al., 2005). Four *TOX* genes are known in *F. graminearum* with *TOX1*, *TOX2*, and *TOX3* being present in a gene cluster while *TOX4* is located at the opposite end of the chromosome. The four genes showed tissue-specific differential expression with *TOX1*, *TOX2*, and *TOX3* being upregulated in FHB while TOX4 was upregulated in FRR (Figure 7B). RT-qPCR demonstrated that the differential expression of *TOX2* was maintained over time (Figure 7C). Expression of *TOX1*, *TOX2*, and *TOX3* was reported to be upregulated in FHB of wheat and barley (Lysøe et al., 2011, Harris et al., 2016, Pan et al., 2018) whereas *TOX4* was only expressed in wheat FHB (Lysøe et al., 2011, Harris et al., 2016, Pan et al., 2018). The role of *TOX* genes in *F. graminearum* virulence in FHB and FRR is unclear but the host/tissue-specific expression of the four genes is intriguing. The examples above hint at the possibility of a specialised secretome where *F. graminearum* has the capacity to recognise the type of host tissue it is infecting and deploy a bespoke array of effectors.

To summarise, the transcriptome response of *B. distachyon* and *F. graminearum* during early FHB and FRR was investigated and compared. There were similarities in *B. distachyon* spike and root tissues with increased expression of genes associated with antimicrobial compounds and JA and ethylene phytohormones-associated genes in both tissues. Significantly, however, there were several tissue-dependent responses including those involved with the phytohormones cytokinin and auxin, receptor signalling, cell-wall modification, and ROS metabolism and response. Likewise for *F. graminearum*, there were both core-genes that were expressed in a tissue-independent manner such as DON and CWDEs, while *TOX* genes, and those involved in cutin degradation, and aurofusarin biosynthesis were expressed in a tissue-specific manner. Overall, this study reveals both host defence responses and pathogen attack strategies in different tissues of the same host and highlights some potentially important tissue-specific aspects.

## Supporting information

Bd RNA-seq Data

Fg RNA-seq Data

## Acknowledgments

This work was supported by the BBSRC (grant number: BB/M011216/1) and BASF SE at Limburgerhof in Germany as part of the PhD studentship of John F. Haidoulis and is supported by the BBSRC Plant Health ISP (grant number: BBS/E/J/000PR9797) for Paul Nicholson.

## Supplemental Material Captions

<u>Supplementary Figure S1:</u> Expression time-course of JA-associated *B. distachyon* genes in FHB and FRR. The blue lines denote FHB and orange lines denote FRR. The Log values presented are calculated by comparing infected tissue against mock-inoculated treatments. The reference housekeeping gene GAPDH was used. Each point is the average of three biological replicates and 2-3 technical replicates. Levels of significance relative to the mock-control, Cq t-test * p<0.05, ** p<0.01, *** p < 0.001.

<u>Supplementary Table S1:</u> Primer Lists for time-course B. distachyon and F. graminearum genes.

<u>Supplementary Table S2:</u> *F. graminearum* functional pathways that are significantly expressed in FHB and FRR from gene-list enrichment analysis.

<u>Supplementary Table S3:</u> *B. distachyon* RNA-seq complete dataset. Gene counts is the average between three biological replicates.

<u>Supplementary Table S4:</u> *B. distachyon* RNA-seq gene sorting per treatment and expression direction.

<u>Supplementary Table S5:</u> *B. distachyon* RNA-seq gene-list enrichment analysis.

<u>Supplementary Table S6:</u> *B. distachyon* RNA-seq phytohormone-related genes.

<u>Supplementary Table S7:</u> *B. distachyon* RNA-seq all genes grouped into putative functional groups.

<u>Supplementary Table S8:</u> *F. graminearum* RNA-seq complete dataset for FHB and in vitro control treatments.

<u>Supplementary Table S9:</u> *F. graminearum* RNA-seq complete dataset for FRR and in vitro control treatments.

<u>Supplementary Table S10:</u> *F. graminearum* RNA-seq gene sorting per treatment and expression direction.

<u>Supplementary Table S11:</u> *F. graminearum* RNA-seq filtered effector genes (Brown et al. 2012) sorted per treatment and expression direction.

<u>Supplementary Table S12.</u> Comparison of hormone pre-treatment and hormone-associated gene expression in FHB and FRR between (Haidoulis and Nicholson, 2020) and this study.

