## Supplemental Figure and Tables for "Tissue-Specific Transcriptome Responses to Fusarium Head Blight and Fusarium Root Rot"

These tables and figure supplement the following manuscript:

By John F Haidoulis and Paul Nicholson

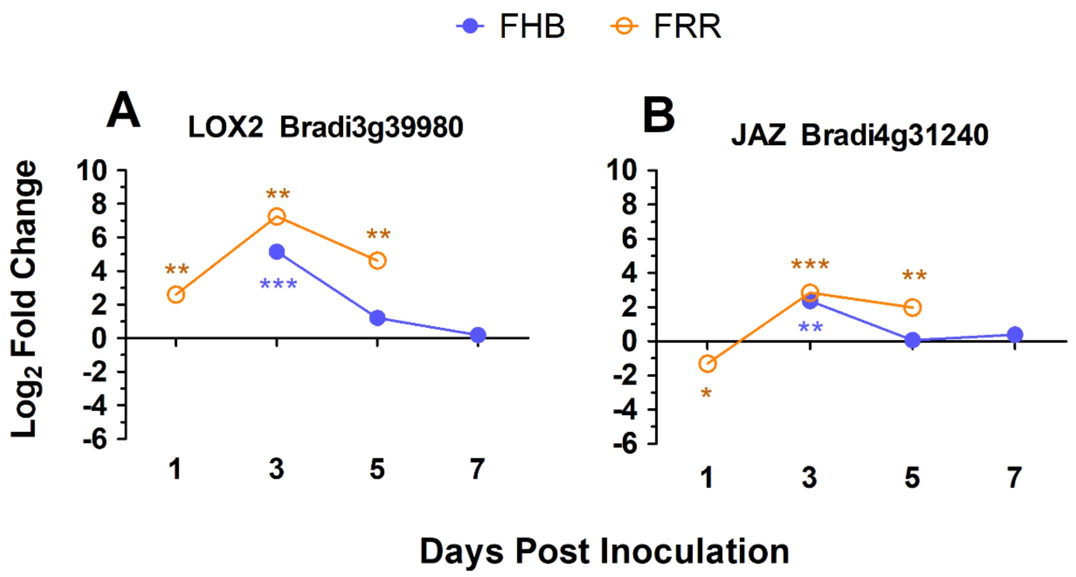


Supplementary Figure S1: Expression time-course of JA-associated *B. distachyon* genes in FHB and FRR. The blue lines denote FHB and orange lines denote FRR. The Log values presented are calculated by comparing infected tissue against mock-inoculated treatments. The reference housekeeping gene GAPDH was used. Each point is the average of three biological replicates and 2-3 technical replicates. Levels of significance relative to the mock-control, Cq t-test * $p<0.05$, ** $p<0.01$, *** $p < 0.001$.

Supplementary Table S1. Primer Lists for time-course *B. distachyon* and *F. graminearum* genes.

| Gene function | Gene name | Gene ID | Gene ID in Figures | Primer Pair (F/R) | Tm (°C) | Product size (bp) |
|---|---|---|---|---|---|---|
| Bd3-1 HK gene | *GAPDH** | DV482924<br>Bradi3g14120 † | *GAPDH* | - TTGCTCTCCAGAGCGATGAC - /<br>- CTCCACGACATAATCGGCAC - | 50-60 | 236 |
| SA/JA | *PR1*** | Bradi1g57590 | *PR1* | - CGAGAAGAAGAACTACCACCATGAC - /<br>- ACACCCGATGGCAGTCGA - | | |
| SA | *NPR4* | Bradi2g54340 | *NPR4* | - ATGGAGTTGCGGTTGTTGTC - /<br>- GCCAGTGATGTGAACAGAGC - | 59.05<br>59.20 | 82 |
| SA | *AtMES1*** | Bradi2g52110 | *MES1* | - AGCTGCCTATTTCCATGCTGTT - /<br>- ATCGAACCCACTCGCATCA - | | |
| SA | *WRKY45_1* | Bradi2g30695 | *WRKY* | - CACAAGTACGACCAGCAGTG - /<br>- GCCGATGTATGTCACCCTGA - | 58.86<br>59.54 | 96 |
| JA | *JAZ* | Bradi4g31240 | *JAZ* | - CGGCAGCTGACCATCTTTTA - /<br>- CTCTGTGCAGGTTGGGGC - | 58.26<br>60.67 | 164 |
| JA | *LOX2* | Bradi3g39980 | *LOX2* | - CCATCGATAAGAGCACACGT - /<br>- GAGGAGGAAGCAAGGACATG - | 57.79<br>57.96 | 185 |
| Auxin | *Aux/IAA* | Bradi1g09090 | *AUXIA* | - CCACCAGTCCGATCGTACC - /<br>- TCCTTCTCGGCTTCCTCTTC - | 59.56<br>58.81 | 80 |
| Auxin | *GH3-2* | Bradi2g50840 | *GH3* | - GTCCCCGTGGTCACCTAC - /<br>- TTGGGTGGGAGGAGATGATG - | 59.02<br>58.78 | 88 |
| CK | *LOG1* | Bradi2g42190 | *LOG1* | - CATCGACCTGGTCTACGGAG - /<br>- CAATCACATGGCGTCCTCC - | 59.34<br>58.6 | 98 |
| CK | *A-type RR 9* | Bradi4g43090 | *RR9* | - CAACAGCTGTAACCCCACAA - /<br>- TGTTGTTTGCAGAGTCGGTG - | 58.31<br>58.99 | 80 |
| Don Detox/resistance | *CYP450* | Bradi2g44150 | *CYP450* | - GCCCGTATTGTTCATCCACC - /<br>- ATTCTTGGACGCCTTGGAGA - | 58.98<br>59.02 | 103 |
| Bd ROS Burst | *Oxalate Oxidase* | Bradi1g11930 | *OXA* | - ATCGTCTTTGTTCCGCTCAC - /<br>- AGGCGAACTTGGACTTGAGA - | 58.57<br>58.95 | 118 |
| ABC transporter (ATPase transporter) | *ABC transporter* | Bradi2g43120 | *ABC* | - CGTGCCGGTTAAGGACTTTG - /<br>- CTCTTGCTCTTGTCGAACGG - | 59.48<br>58.94 | 96 |
| Fungal HK gene | ubiquitin C-teminal hydrolase*** | FGSG_01231.3 | *gzUBH* | -GTTCTCGAGGCCAGCAAAAAGTCA - /<br>- CGAATCGCCGTTAGGGGTGTCTG - | 62.4<br>65.2 | 168 |
| Fungal Predicted Effector | *TOX2* | FGRAMPH1_01G00199 | *Tox2* | - CTACAGGCCCTTCTTGACCA - /<br>- GTCAACTGCCCATCATCGAC - | 59.02<br>58.99 | 86 |
| Fungal Predicted Effector | Pectate Lyase | FGRAMPH1_01G16515 | PecLy | -GTACCAAGACCCTCAGCACT - /<br>- AGACTGGCCGGTACATTTCA - | 59.02<br>59.02 | 101 |

Primer pairs were designed on Primer 3 (Kõressaar et al., 2018, Untergasser et al., 2012, Koressaar and Remm, 2007) unless otherwise stated: * From (Hong et al., 2008). ** From (Kakei et al., 2015). *** From (Kim and Yun, 2011). † Predicted as this geneID on (Ensembl Genomes database (Howe et al., 2020)). Abbreviation: HK (Housekeeping), F (Forward primer), R (Reverse primer). Hormone abbreviations: SA (Salicylic acid), JA (Jasmonic acid), CK (Cytokinin).

Supplementary Table S2. *F. graminearum* functional pathways that are significantly expressed in FHB and FRR from gene-list enrichment analysis.

| Pathway | KEGG Pathway ID | Number of genes | p-adj |
|---|---|---|---|
| **Upregulated in response to FHB** | | | |
| Metabolic pathways | fgr01100 | 204 | p < 0.001 |
| Pentose and glucuronate interconversion | fgr00040 | 22 | p < 0.001 |
| Starch and sucrose metabolism | fgr00500 | 22 | p < 0.001 |
| Tyrosine metabolism | fgr00350 | 22 | p < 0.001 |
| Valine, leucine, and isoleucine degradation | fgr00280 | 19 | p < 0.001 |
| Cyanoamino acid metabolism | fgr00460 | 13 | p < 0.01 |
| Other glycan degradation | fgr00511 | 8 | p < 0.01 |
| Beta-Alanine metabolism | fgr00410 | 13 | p < 0.01 |
| Biosynthesis of secondary metabolites | fgr01110 | 65 | p < 0.01 |
| **Upregulated in response to FRR** | | | |
| Metabolic pathways | fgr01100 | 178 | p < 0.001 |
| Valine, leucine, and isoleucine degradation | fgr00280 | 21 | p < 0.001 |
| Tyrosine metabolism | fgr00350 | 19 | p < 0.01 |
| Beta-Alanine metabolism | fgr00410 | 14 | p < 0.01 |
| Pentose and glucuronate interconversion | fgr00040 | 13 | p < 0.05 |
| Starch and sucrose metabolism | fgr00500 | 15 | p < 0.05 |
| Biosynthesis of secondary metabolites | fgr01110 | 60 | p < 0.05 |
| Arginine and proline metabolism | fgr00330 | 14 | p < 0.05 |

Based on output from KOBAS 3.0 web-based browser software.

Tables S3-S7: RNA-seq *B. distachyon* data tables.

- S3: *B. distachyon* RNA-seq complete dataset. Gene counts is the average between three biological replicates.
- S4: *B. distachyon* RNA-seq gene sorting per treatment and expression direction.
- S5: *B. distachyon* RNA-seq gene-list enrichment analysis.
- S6: *B. distachyon* RNA-seq phytohormone-related genes.
- S7: *B. distachyon* RNA-seq all genes grouped into putative functional groups.

Tables S8-S12: RNA-seq *F. graminearum* data tables.

- S8: *F. graminearum* RNA-seq complete dataset for FHB and in vitro control treatments.
- S9: *F. graminearum* RNA-seq complete dataset for FRR and in vitro control treatments.
- S10: *F. graminearum* RNA-seq gene sorting per treatment and expression direction.
- S11: *F. graminearum* RNA-seq filtered effector genes (Brown et al. 2012) sorted per treatment and expression direction.

Supplementary Table S12. Comparison of hormone pre-treatment and hormone-associated gene expression in FHB and FRR between (Haidoulis and Nicholson, 2020) and this study.

| | **Hormone pre-treatment (Haidoulis and Nicholson, 2020)** | | | | **Gene expression (This study)** | | |
|---|---|---|---|---|---|---|---|
| Hormone | FHB | FRR | Tissue effect | | FHB | FRR | Tissue effect |
| JA | Susceptible | Resistant | opposite | | Up | Up | same |
| ET | Susceptible | Resistant | opposite | | Up | Up | same |
| SA | None | Susceptible | - | | - | - | - |
| Auxin | Resistant | Resistant | same | | Up | Down | opposite |
| Cytokinin | Susceptible | Susceptible | same | | Up | Down | opposite |
